# An enhancer-centric approach applied to human immune system epigenomes revealed the association of macrophage enhancers with cardiovascular disease

**DOI:** 10.64898/2026.06.02.728745

**Authors:** Juliana E Arcila-Galvis, Felipe Were, David Juan, Jose Del-Rio, Christopher A. Lamb, Sophie Hambleton, Antonio Castrillo, Salvatore Spicuglia, Alfonso Valencia, Ines Pineda-Torra, Enrique Carrillo-de-Santa-Pau, Robert P. Hirt, Daniel Rico

## Abstract

Complex diseases are influenced by both genetic and environmental factors. Immune cells are key mediating interactions with the environment, but the impact of genetic variation on the immune system and how it influences complex diseases is not fully understood. Moreover, most genome-wide analyses (GWAS) variants associated with complex diseases are non-coding and difficult to interpret. Here, we investigated the association of non-coding variants with immune cell enhancers. As part of BLUEPRINT and the International Human Epigenome (IHEC) consortia, we generated and analysed a comprehensive set of epigenomes for human primary immune cells, including 107 epigenomes derived from 749 ChIP-Seq experiments across 24 cell types. We identified multicell enhancer activity patterns across the genome and examined their links with non-coding variants from 518 GWAS traits. This analysis revealed 117 significant associations, including novel links between cardiovascular disease variants and macrophage-specific enhancers that regulate genes involved in lipid metabolism and immunity, such as the gene encoding for the nuclear receptor LXR-alpha (*NR1H3)* and many of its known target genes. Together, these data will help to better understand the influence of genetic variability in immune function and related diseases.

## Introduction

The understanding of inherited diseases has been historically driven by the discovery of highly penetrant pathogenic variants in affected family members [1], which provide a clear mechanistic link to disease by altering the function or expression of a specific protein. Most human traits, however, result from complex interactions between non-coding genotypes at multiple loci and environmental cues [2], involving multiple biological processes acting across temporal and spatial scales [3]. Consequently, the mechanistic coherence between genomic variants and phenotypes seen in Mendelian diseases is largely lacking in complex diseases [3]

For a multitude of complex traits, genome-wide association studies (GWAS) have identified numerous widespread genomic variants, predominantly non-coding, that weakly influence these traits [4,5]; however, the pervasive lack of understanding of how these variants interact to establish disease mechanisms has been referred to as the GWAS coherence problem [3]. Indeed, GWAS studies alone typically fail to provide substantial evidence to guide functional genetic analysis experimentation for many traits [6].

There have been significant advances in addressing the coherence problem, particularly through the integration of transcriptomic and epigenomic data to link non-coding variants to their function [7]. It is well established that disruption of enhancers and other non-coding regulatory elements can lead to disease by perturbing the fine-tuning of gene expression [8]. Accordingly, previous studies have integrated GWAS signals with large collections of epigenomic maps to infer the cell types in which non-coding disease-associated variants may drive pathogenic phenotypes [9–16] Variant enrichment approaches, such as linkage disequilibrium (LD), score regression and fine-mapping, have further enabled the systematic linking of GWAS signals to regulatory annotations[17,18].

Despite the undeniable advances these studies have brought to our understanding of genotype–phenotype interactions in complex diseases, they have not fully bridged the coherence gap, in part due to their simplistic modelling of gene regulatory processes. Variant-centric methods typically treat enhancers as isolated units, whereas in reality, regulatory elements often act in coordinated clusters across cell types [19,20] The set of variants associated with a disease can therefore span clusters of enhancers or extended regulatory regions, which tend to have stronger effects on gene expression than single elements [21–25] and are enriched in transcription factor binding sites, often harbouring multiple causal variants [24,26,27]. Consistent with this, enhancer proximity has been shown to produce super-additive effects on gene expression, whereby perturbation of one enhancer influences neighbouring regulatory elements [28]. In this context, modelling a single variant within a single cell type is unlikely to capture its full physiological consequences [29].

Moreover, epigenomic maps used in these analyses, such as those generated by the ENCODE and NIH Roadmap Epigenomics consortia [15,30], have a limited representation of immune cells. Yet these cells are central to interpreting non-coding variation in complex traits, as disease onset is strongly shaped by environmental factors and inflammatory cues, and immune cells are key mediators of responses to these stimuli through highly dynamic and complex enhancer landscapes that enable rapid modulation of gene expression, central to the coordination of myriad of processes underlying effective immune responses [31,32]. To fill this gap, the BLUEPRINT Consortium was funded by the European Union to generate reference epigenomes of haematopoietic cell types in healthy humans, including most immune cell types in the lymphoid and myeloid lineages [33]. BLUEPRINT was part of the wider International Human Epigenome Consortium (IHEC) that coordinates standardisation of protocols across participating projects, recently also including data reprocessing and metadata harmonisation efforts through the IHEC EpiAtlas initiative (International Human Epigenome Consortium, in preparation).

Here, we generated 107 epigenomic maps across 31 primary haematopoietic cell types using data from the BLUEPRINT Consortium [33], representing the largest and most comprehensive collection of reference haematopoietic epigenomes to date and that were not included in previous studies integrating GWAS signals and enhancers. We generated a collection of 1,100 associations between sets of enhancers active in immune cells and more than 172 complex traits, including cardiovascular, cognitive, cancer, and autoimmune traits. Additionally, to establish connections between enhancers and traits, we employed an enhancer-centric approach, identifying broader functionally relevant regions associated with disease and considering their multicellular activity profiles.

Additionally, by analysing groups of variants that act in the same subset of cell types, we can generate a more coherent interpretation of GWAS results and propose disease mechanisms. For instance, we found multiple variants across several macrophage enhancers that seem to modulate cardiovascular disease risk by regulating the expression of genes with key roles in the three main pathways of lipid metabolism. Macrophages are central regulators of lipid handling, inflammation and foam-cell formation in atherosclerosis, highlighting them as a major biologically coherent context in which to interpret non-coding cardiovascular disease (CVD) risk variants [34,35]. Our data are consistent with inferences made from highly penetrant oligogenic forms of dyslipidaemia that predispose to early onset cardiovascular disease [36].

## Results

### Generation of 107 primary human haematopoietic epigenomes and 31 consensus cell type epigenomes

To generate a consistent functional annotation of regulatory elements and transcriptional states across haematopoietic cell types, we generated genome-wide integrated epigenomic maps using histone mark profiles from primary human samples generated by the BLUEPRINT Consortium [33] (**Table S1**). Our dataset comprises 107 reference epigenomes spanning 31 cell types. These include multiple lymphoid and myeloid populations, such as five T-cell subtypes, six B-cell subtypes, natural killer cells, mature neutrophils and four of their progenitor states, as well as monocytes and six additional monocyte-differentiated cell types. Mesenchymal stem cells and endothelial cell types were also included to provide a broader cellular context (**Fig. 1A–C).**

**Figure 1.**
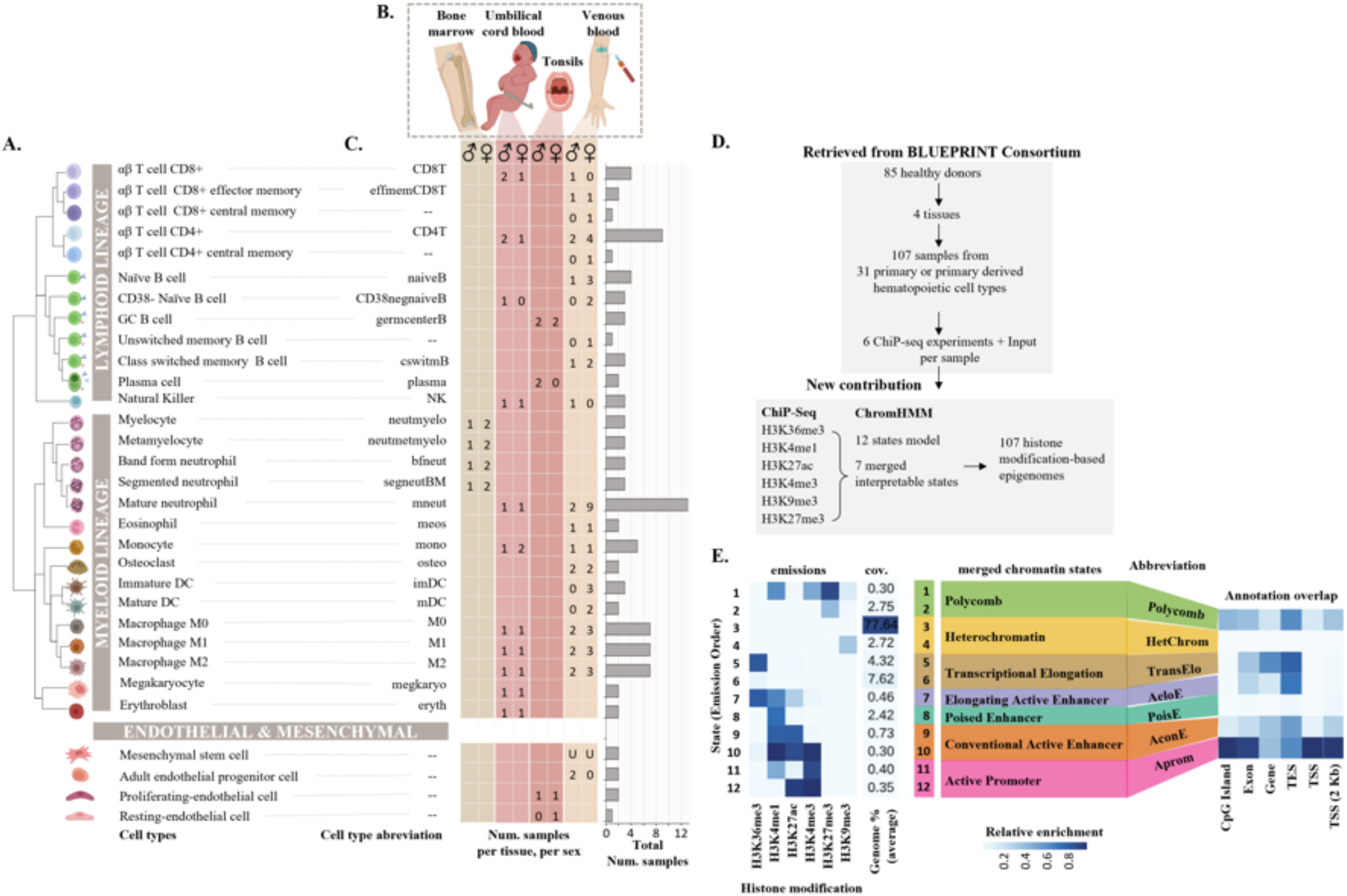
Dataset description. **(A)** Hematopoietic cell types included in the analysis and their lineage hierarchy. **(B)** Tissues from which the cells were sampled **(C)** Counts of biological replicate samples for each cell type, tissue, and sex. **(D)** A simplified diagram describing the BLUEPRINT Consortium data used to derive the 107 epigenomes newly generated in this study, including ChIP-seq profiles of six histone modifications and matched controls for each sample **(E)** The heatmap in the left panel displays the ChromHMM emission probabilities for the 12-state model generated in this study. The middle panel shows the state annotations based on literature and genomic annotation enrichments. The right panel shows genomic annotation enrichments for each state after collapsing the 12-state model into seven interpretable chromatin states. Abbreviations: TSS, transcription start site; TES, transcription end site.

The BLUEPRINT dataset comprises 642 independent ChIP–seq experiments profiling six histone marks (H3K4me3, H3K4me1, H3K27ac, H3K27me3, H3K36me3 and H3K9me3) together with input controls in each sample, enabling complete chromatin state annotation without the need for imputation. Using these data, we trained a ChromHMM model [37] to identify combinatorial patterns of histone marks across the genome (**Fig. 1D**). This analysis yielded 12 chromatin states consistent with previously described models from the NIH Roadmap Epigenomics Consortium [15] (**Fig. S1, Tables S2 and S3)**.

For interpretability, these were grouped into seven categories reflecting regulatory and transcriptional functions, including: active promoter (Aprom), transcriptional elongation (TransElo), Polycomb, heterochromatin (HetChrom), conventional active enhancers (AconE), active elongating enhancers (AeloE), and poised enhancers (PoiseE) (**Fig. 1E**).

To enable the comparison of epigenomes across cell types, we generated a consensus epigenome for each cell type with at least two biological replicates (27 out of 31 cell types; **Fig. 1A**). Consensus states were defined as chromatin annotations consistently observed in at least 75% of replicates. In addition to the seven chromatin states, we introduced a ‘non-conserved’ state to capture genomic regions lacking consistent annotation across replicates (**Fig. 2A)**.

**Figure 2.**
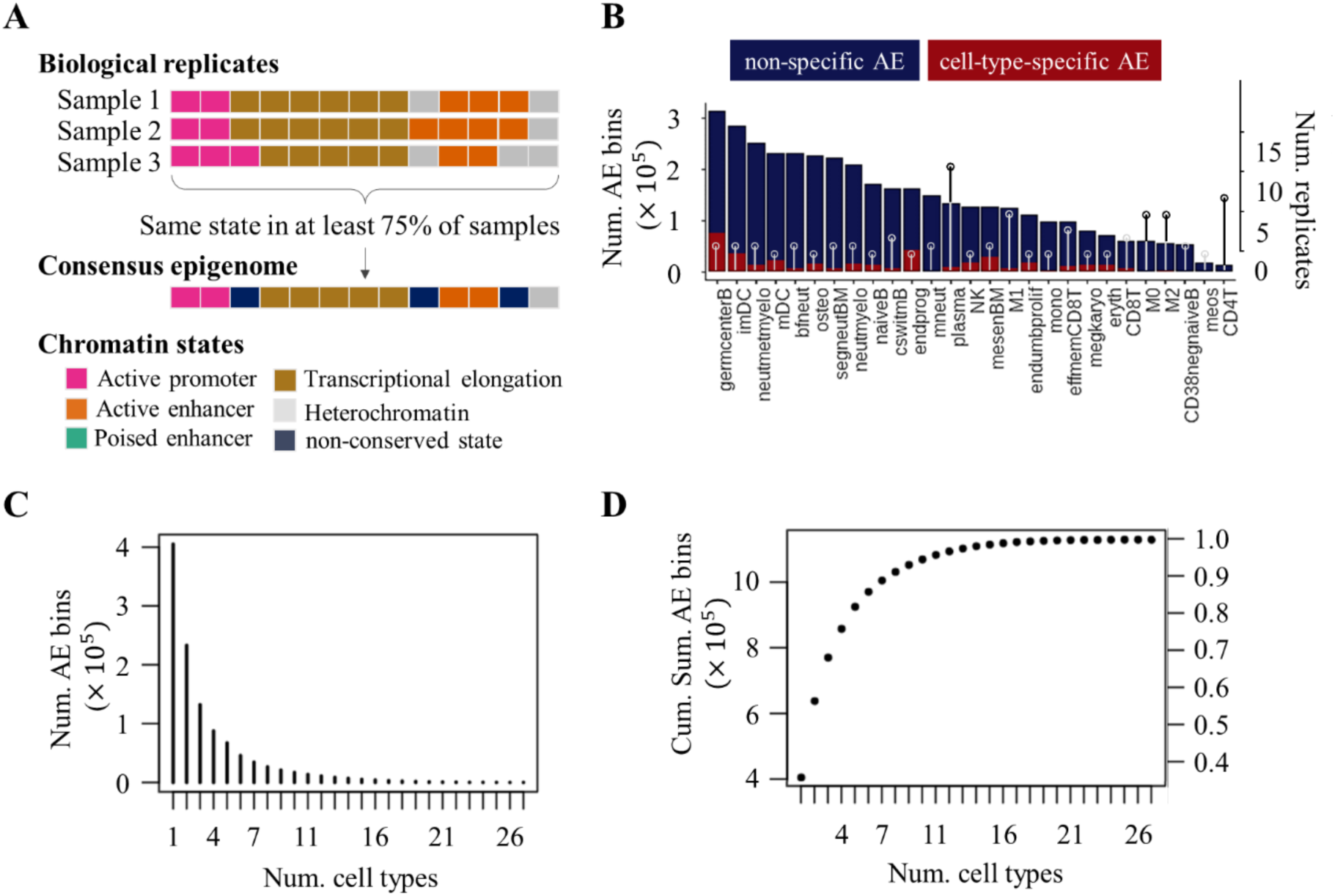
Cell type specificity and sharing of active enhancers across consensus epigenomes. **(A)** For each cell type, a consensus epigenome was generated by assigning the state conserved in at least 75% of the samples available for that cell type. The figure shows chromatin state annotations for a hypothetical genome fragment. Each rectangle represents a200 bp bin annotated using a chromatin states model. **(B)** Bar plot showing the number of AE bins (left y-axis) in each cell type’s consensus epigenome. Red bars indicate cell type-specific enhancers, while blue bars represent enhancers shared across cell types. The number of biological replicates (right y-axis) for each cell type is shown as lollipops. (C) Frequency plot displaying the number of AE bins (y-axis) unique to a single cell type (x=1) or shared across two or more cell types (2 ≤ x ≤ 27). (D) Cumulative count of AE bins (y-axis) and their equivalent percentage of the total consensus AE categorized as either unique to a single cell type (x=1) or shared among multiple cell types (2 ≤ x ≤ 27).

Across all cell types, 46% of the genome was consistently assigned the same consensus chromatin state across all cell types, predominantly heterochromatin, whereas the remaining genomic regions displayed variability, highlighting the dynamic nature of regulatory landscapes (**Fig. S2, top panel**).

We next examined the proportion of each consensus epigenome covered by active enhancer regions (AE: AeloE and AconE). Most enhancers active in a given cell type were also active in at least one additional cell type (**Fig. 2B–C**), with on average only 10% being cell type-specific (range: 1.3%–26.4%; **Fig. 2B**). When aggregated across all cell types, however, these cell type-specific enhancers accounted for 40% of the total haematopoietic enhancer repertoire (**Fig. 2C–D, Table S4**).

The number of AE bins (200 bp genomic segments) varied widely across cell types, ranging from 15,268 to 313,787 (mean 143,300; **Table S4**). Germinal centre B cells (germcenterB; 313,787 bins) and immature dendritic cells (imDC; 285,986 bins) exhibited the highest numbers of AE bins, whereas CD4+ T cells (CD4T; 15,268 bins) and mature eosinophils (meos; 17,626 bins) showed markedly fewer enhancers (**Fig. 2B**).

To assess whether this variability reflects biological rather than technical differences, we examined the relationship between the number of consensus AE bins per cell type and the number of biological replicates, observing only a weak correlation (Pearson r = 0.3). In contrast, the number of consensus enhancers showed a strong correlation with the mean number of enhancers per cell type (Pearson r = 0.82), suggesting that the reduced enhancer counts in eosinophils and CD4+ T cells are unlikely to be driven by replicate number and may instead reflect underlying biological differences.

Finally, we calculated the cumulative number of AE bins as a function of the number of cell types in which they are active (**Fig. 2C-D**). The number of AE bins increased rapidly with the inclusion of additional cell types before reaching a plateau, suggesting near-saturation of enhancer discovery across the major haematopoietic cell types in resting/baseline state, while acknowledging that context-specific regulatory elements may remain uncharacterised.

### Defining enhancer activity profiles across haematopoietic cell types

Given that the same genomic region can adopt distinct regulatory states across cell types [9,38], we defined enhancer activity profiles to capture the multi-cell-type activity of enhancers. Each profile corresponds to the set of cell types in which a genomic region is consistently annotated as an active enhancer in the consensus epigenomes (**Fig. 3A–B**). Across all genomic regions annotated as active enhancers in at least one of the 27 cell types with biological replicates, we identified 107,562 distinct activity profiles, reflecting the high combinatorial diversity of enhancer activity.

**Figure 3.**
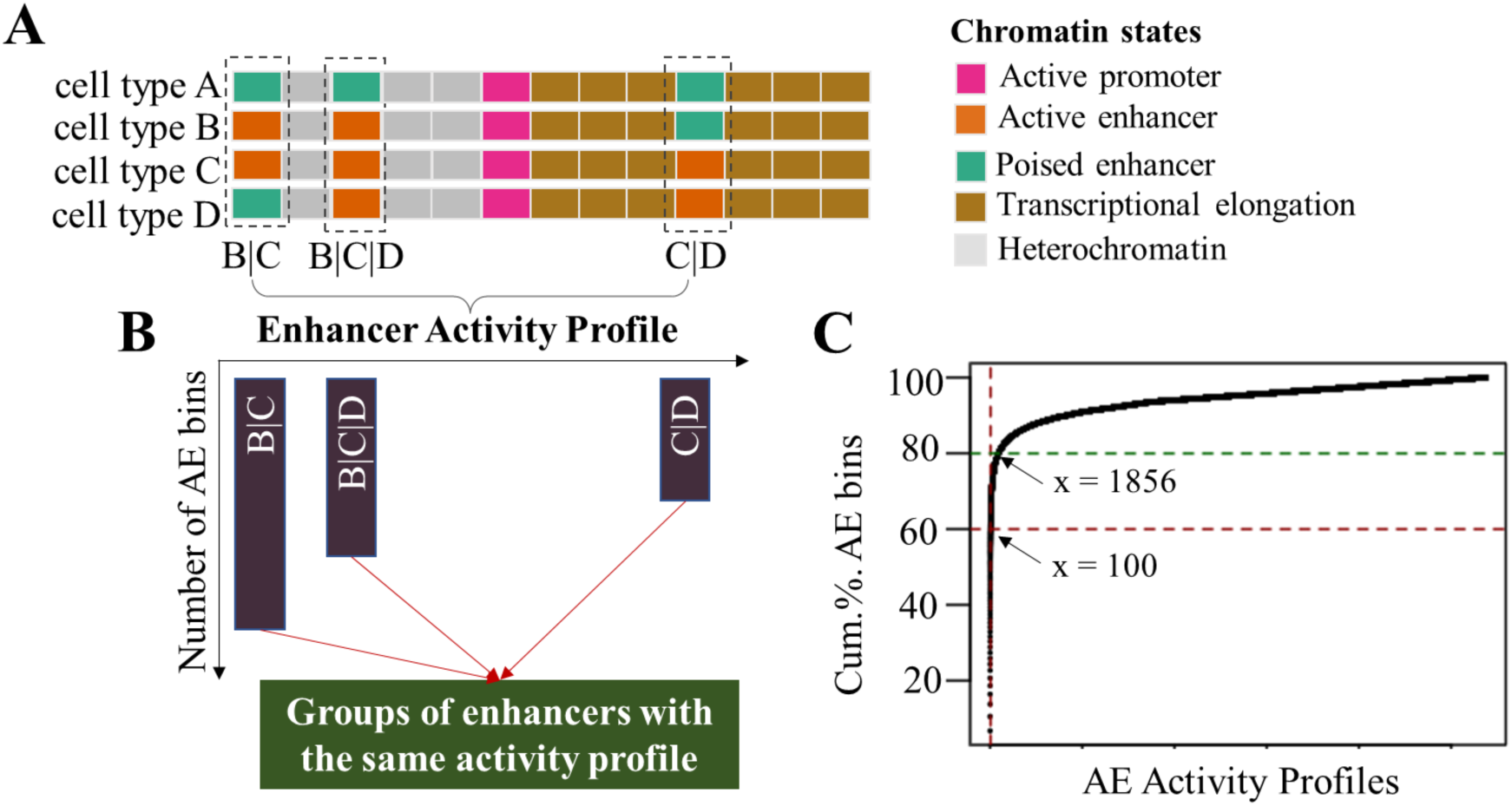
Enhancer activity profiles and their prevalence in haematopoietic cells. **(A)** Enhancer activity profiles were assigned to each active enhancer (AE) bin across the genome based on the set of cell types in which the enhancer was annotated as an active enhancer. The figure depicts hypothetical chromatin state annotations for a genome fragment. Each coloured rectangle represents a 200 bp bin annotated using the chromatin states model. Bins enclosed in dashed black rectangles contain enhancers, and their activity profiles are indicated at the bottom using cell type names separated by pipes (“|”). **(B)** Enhancers were grouped based on their activity profiles, with the size of each group measured by the number of 200 bp bins it contains. This reflects how frequently each activity profile appears across the genome in the total set of analysed epigenomes, as some profiles are more common than others. **(C)** Cumulative frequency of enhancer activity profiles: The top 100 most common activity profiles account for 60% of all AE bins. Remarkably, 1,856 profiles make up 80% of all active enhancers, showing that a small number of profiles are responsible for the majority of enhancer activity in the genome among the considered cell types

The frequency of enhancer activity profiles was highly uneven, with most profiles occurring in only a small number of genomic bins (mean 10.5; Q1 = 1; Q3 = 2; range 1–75,814), while a relatively small number of recurrent multi-cell-type patterns accounted for a large fraction of enhancer activity. The 1,856 most frequent profiles accounted for 80% of AE bins, and the top 100 accounted for 60% (**Fig. 3C**).

These top 100 activity profiles were each represented across a substantial number of genomic regions (≥900 AE bins) (**Fig. 4**). The most frequent profile corresponded to cell type-specific enhancer activity in germinal centre B cells (germcenterB; 75,814 bins, 15.16 Mb), these 100 profiles collectively captured cell type specific enhancer activity for the majority of haematopoietic cell types in our dataset (23 out of 27), with the exception of CD38− B cells (CD38negB), inflammatory macrophages (M0), mature eosinophils (meos), and CD4+ T cells. Consistent with their low number of active enhancer regions, neither CD4+ T cells nor mature eosinophils were represented among the most frequent profiles (**Fig. 2B**).

**Figure 4.**
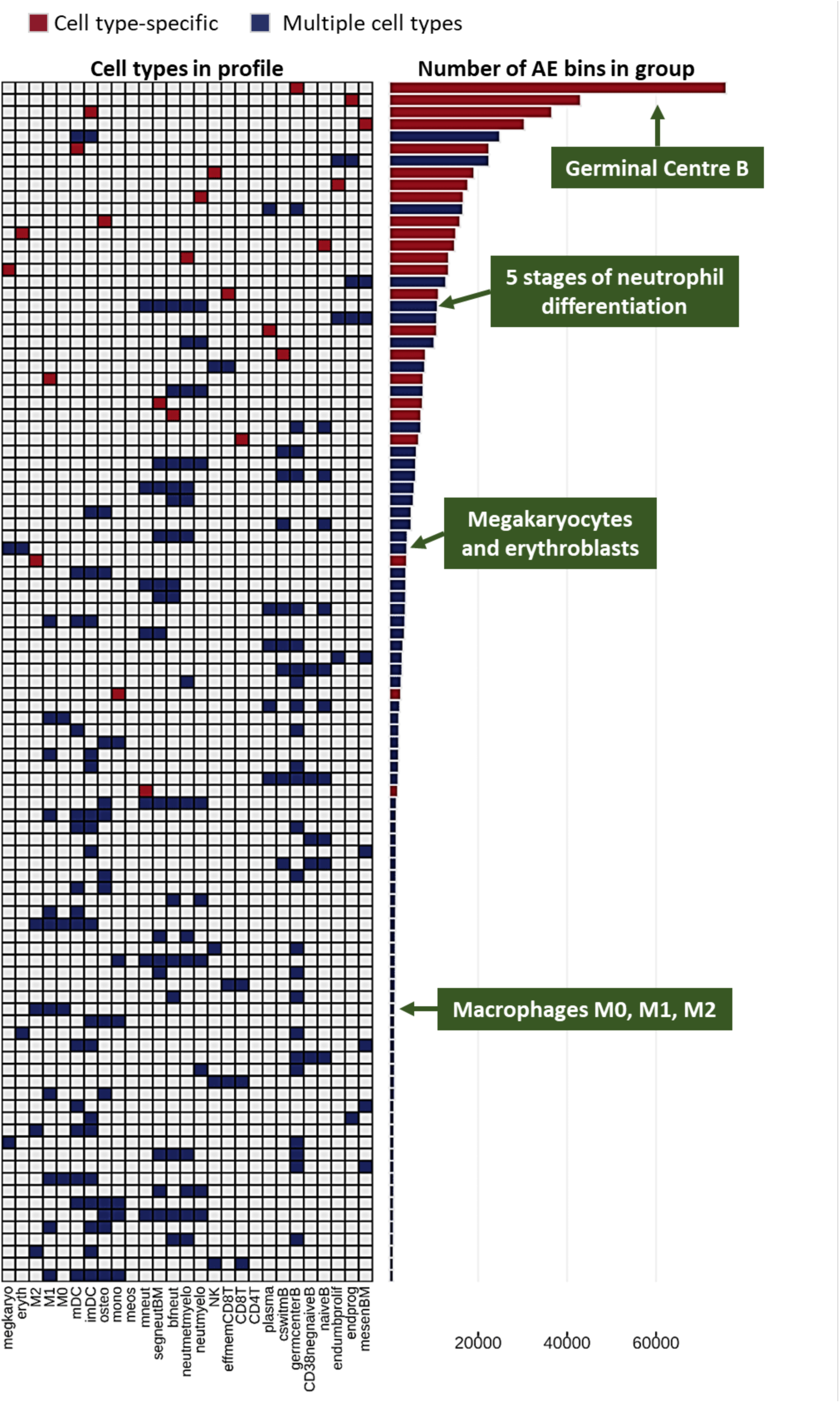
The 100 most frequently observed activity profiles in hematopoietic AE. The heatmap displays the cell types involved in each profile, while the bar plots illustrate the number of AE bins in which each profile was identified. Profiles shown in blue represent AE active in multiple cell types, whereas those in red are cell type-specific AE.

Among these top 100 most common activity profiles, the majority reflected lineage-specific patterns. The profile involving the greatest number of cell types corresponded to enhancers active across seven myeloid populations: monocytes (mono), osteoclasts (osteo), mature neutrophils (mneut) and four neutrophil progenitors (segneutBM, neutmyelo, bfneut and neutmyelo), and encompassed 1,025 genomic bins (205 kb). Similarly, we identified 2,045 bins (409 kb) with enhancer activity across the five stages of B-cell differentiation, and 1,228 bins (245 kb) shared among natural killer (NK) cells, CD8+ T cells (CD8T) and CD8+ effector memory T cells (effmemCD8T) **(Fig. 4).**

Overall, activity profiles involving multiple haematopoietic cell types tended to involve cell types from the same lineage, either lymphoid or myeloid. A small number of profiles deviated from this pattern, comprising enhancers active in both germinal centre B cells and selected myeloid populations, including neutrophil progenitors, monocyte-derived dendritic cells, osteoclasts, megakaryocytes and erythroblasts. This could be partially explained by the fact that germcenterB cells have the consensus epigenome with the highest number of active enhancer regions.

### The activity profiles of GWAS trait-associated enhancers point to relevant cell types in their aetiology

We used these multi-cell-type regulatory patterns to inform the cellular context of non-coding genetic variation associated with complex traits, to that end we integrated haematopoietic enhancer activity profiles with non-coding variants associated with 518 complex traits (**Table S5**) from the GWAS Catalog [4], and assessed the enrichment of active enhancer (AE) bins from the 100 most frequent activity profiles within genomic regions surrounding trait-associated variants (**Fig. 5A**).

**Figure 5.**
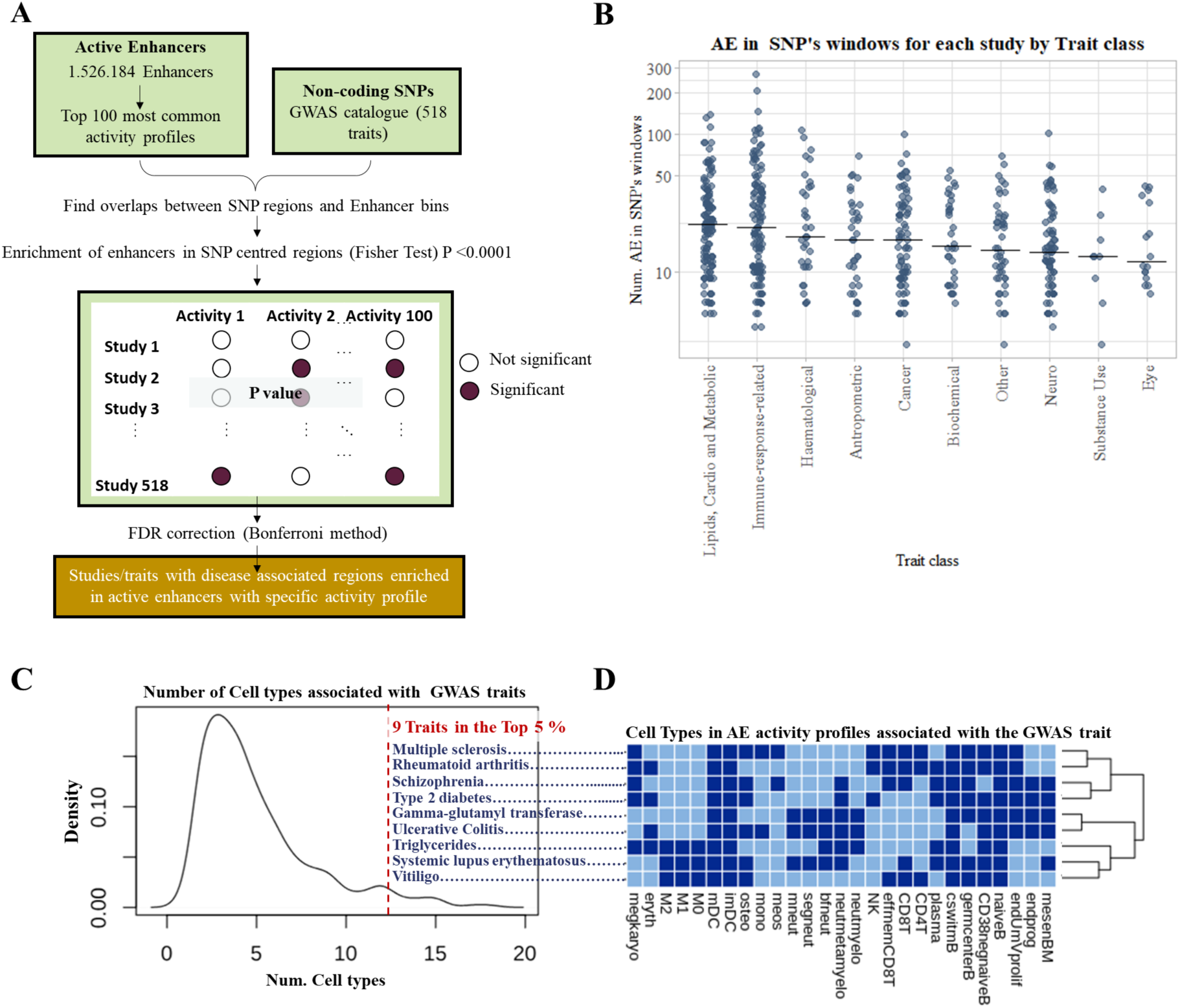
Enhancer-centric enrichment analysis to assess the association between AE activity profiles and common complex diseases. **(A)** Flow diagram illustrating the methodology used to identify trait-associated regions enriched with active enhancer bins, focusing on those with activity profiles among the top 100 most common in our dataset. **(B)** Scatter plot showing the number of AE bins overlapping trait-associated loci for each trait category. Only enhancers with activity profiles significantly associated with the traits in the category are included in the plot. **(C)** Density plot showing the distribution of the number of cell types associated with GWAS traits. The figure illustrates that a single GWAS trait can be associated with enhancers active across various cell types. **(D)** Displays the cell types in AE activity profiles associated with the top 5% GWAS traits from the analysis on panel C. The red dotted line indicates the density point until 95% of the traits are covered. The number of traits within the upper 15% is highlighted in bold red font, emphasising the subset of traits with the most extreme values in bold blue font.

Enrichment was evaluated across 10 kb windows centred on 11,060 non-coding GWAS lead SNPs from 985 studies. This analysis uncovered 1,113 significant associations involving 172 of the 518 traits analysed (**Table S6**). Most traits were not associated with a single regulatory context but instead mapped to multiple enhancer activity profiles across several cell types (typically up to 8 profiles involving up to 12 cell types; **Fig. S3**), indicating that genetic risk is distributed across coordinated regulatory programmes.

To characterise these associations, we grouped traits into 11 biological categories, including lipids, cardiovascular and metabolic traits, immune response-related traits, haematological traits, anthropometric traits, cancer-related traits, biochemical analyses-related traits, eye-related traits, neurological traits, substance use-related traits, and other traits (**Fig. 5B**, **Fig. S4** and **Table S7**).

Traits related to lipids cardiac and metabolic functions displayed the highest median number of AE bins overlapping GWAS loci (**Fig. 5B**). While associations with immune and haematological traits were expected, we also observed substantial enrichment across non-immune traits, including metabolic, cardiovascular and neurological phenotypes, suggesting broader implications of genetic variants in immune cell enhancers.

The cell types contributing to these associations are shown in **Fig. S4**. For example, multiple stages of B-cell differentiation, dendritic cells and effector memory CD8+ T cells (effmemCD8T) were prominently associated with autoimmune conditions **(Fig. S4A)**.

Interestingly, closely related traits, such as ulcerative colitis and Crohn’s disease, did not cluster by enhancer activity profiles, indicating differences in their regulatory architectures **(Fig. S4A)**.

We next examined the number of cell types associated with each trait. Most traits were linked to enhancer activity in a limited number of cell types, typically between two and five (**Fig. 5C**). However, a subset of traits showed associations across a broader range of cell types, forming a long tail in the distribution. This included type 2 diabetes, schizophrenia and several autoimmune diseases such as multiple sclerosis, rheumatoid arthritis, ulcerative colitis, systemic lupus erythematosus and vitiligo. Additional traits associated with enhancer activity across many cell types included gamma-glutamyl transferase and triglyceride levels, markers of hepatic function and cardiovascular disease (CVD) risk, respectively (**Fig. 5D**).

### Most haematopoietic enhancers linked to cardiovascular disease are macrophage-related and colocalise with key players in lipid metabolism

The analysis described above identified more than a thousand enhancer–trait associations across diverse phenotypes. To investigate how these associations translate into disease-relevant regulatory mechanisms, we focused on cardiovascular disease (CVD)-related traits as a representative case, given their substantial contribution to global mortality (∼32%) [39] and their presence in the category with the highest number of enhancer-trait associations (**Fig. 5B**).

CVD-related traits, including levels of LDL cholesterol, HDL cholesterol and triglycerides in the blood, metabolic syndrome, coronary artery disease, waist-hip ratio, and various other lipid-related characteristics, showed widespread enrichment across 133 loci associated with 18 distinct enhancer activity profiles (**Fig. 6**). A substantial proportion of these loci (69 loci, 52%) were linked to macrophage enhancer activity profiles, including unpolarised M0, inflammatory M1 and anti-inflammatory M2 macrophages, as well as combinations thereof (**Fig. 6**). These regions were distributed across 14 chromosomes and could be consolidated into 17 genomic regions based on proximity (**Table 1, Table S8, Fig. S5**).

**Figure 6.**
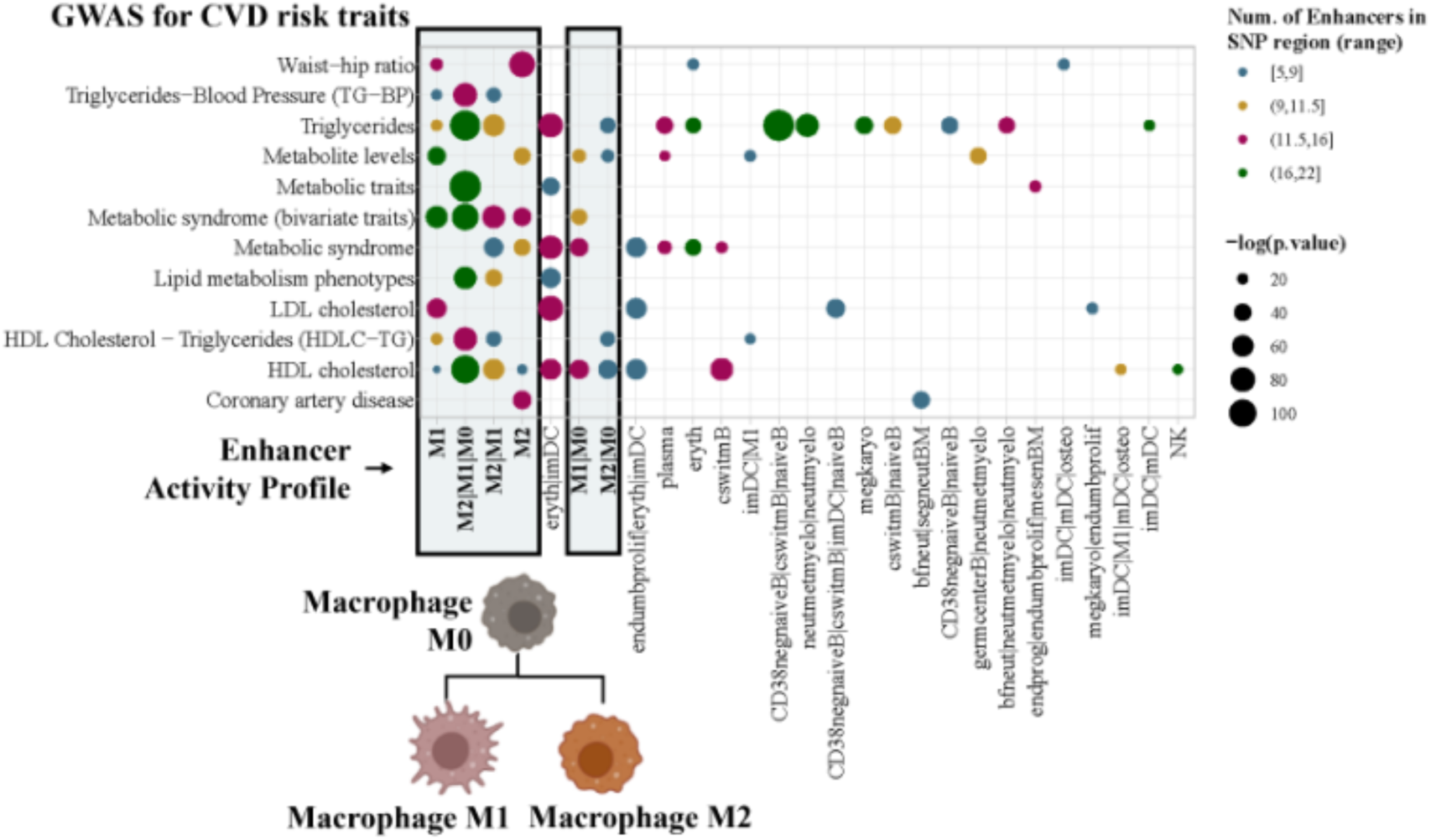
AE activity profiles associated with cardiovascular disease (CVD) risk. Dot plot illustrating enhancer activity profiles enriched in loci associated with CVD risk-related traits. Each dot represents a significant enrichment (padj < 0.0001) observed in at least one GWAS study for the corresponding trait and activity profile. Dot colour represents the minimum number of AE bins overlapping trait-associated loci across GWAS studies, while dot size indicates the significance level (max p adjusted value). Macrophage-specific enhancers, highlighted in blue boxes, show the highest significance and AE bin counts, accounting for 52% of CVD-associated loci. A schematic of macrophage lineage is provided in the bottom left corner.

**Table 1.**
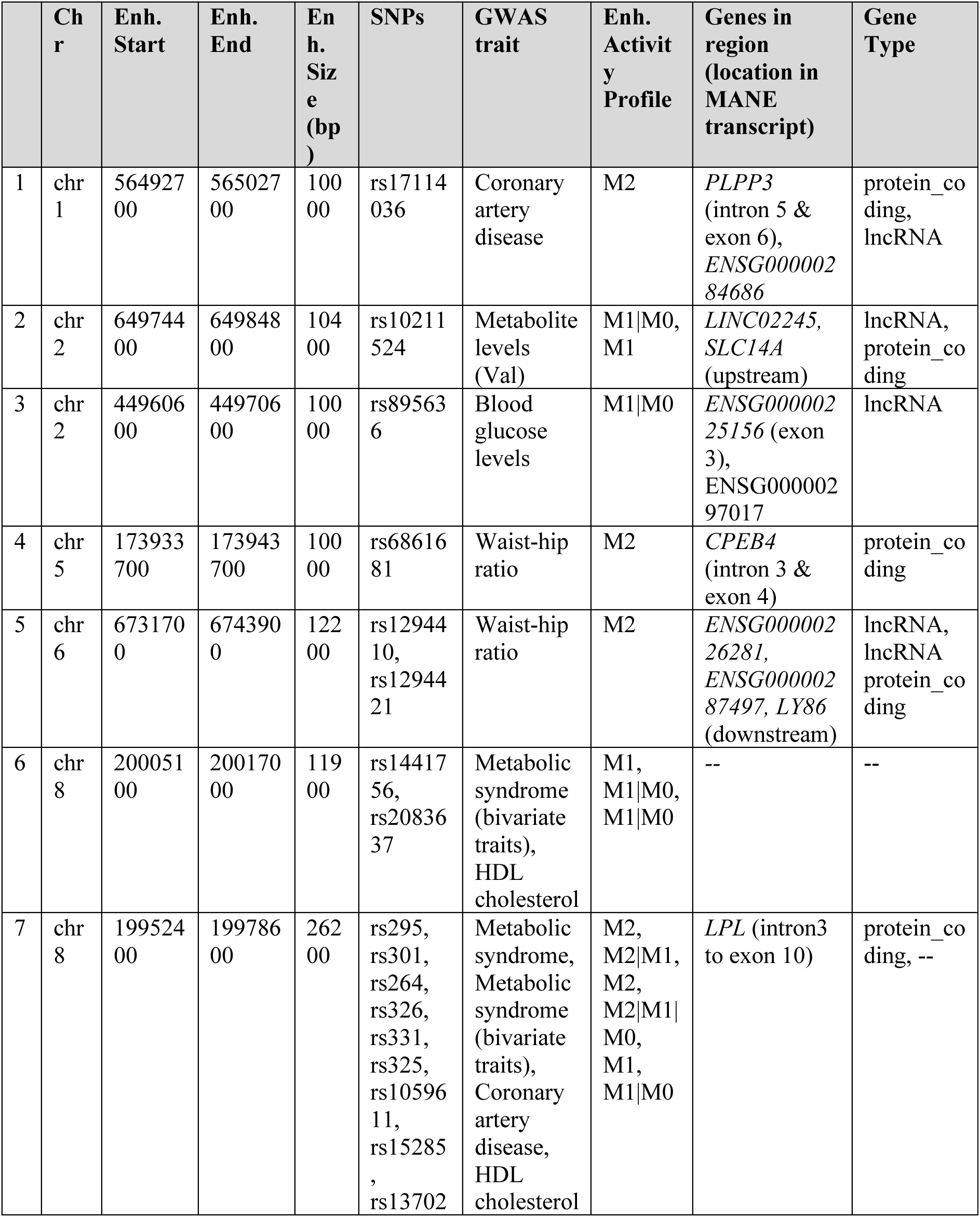
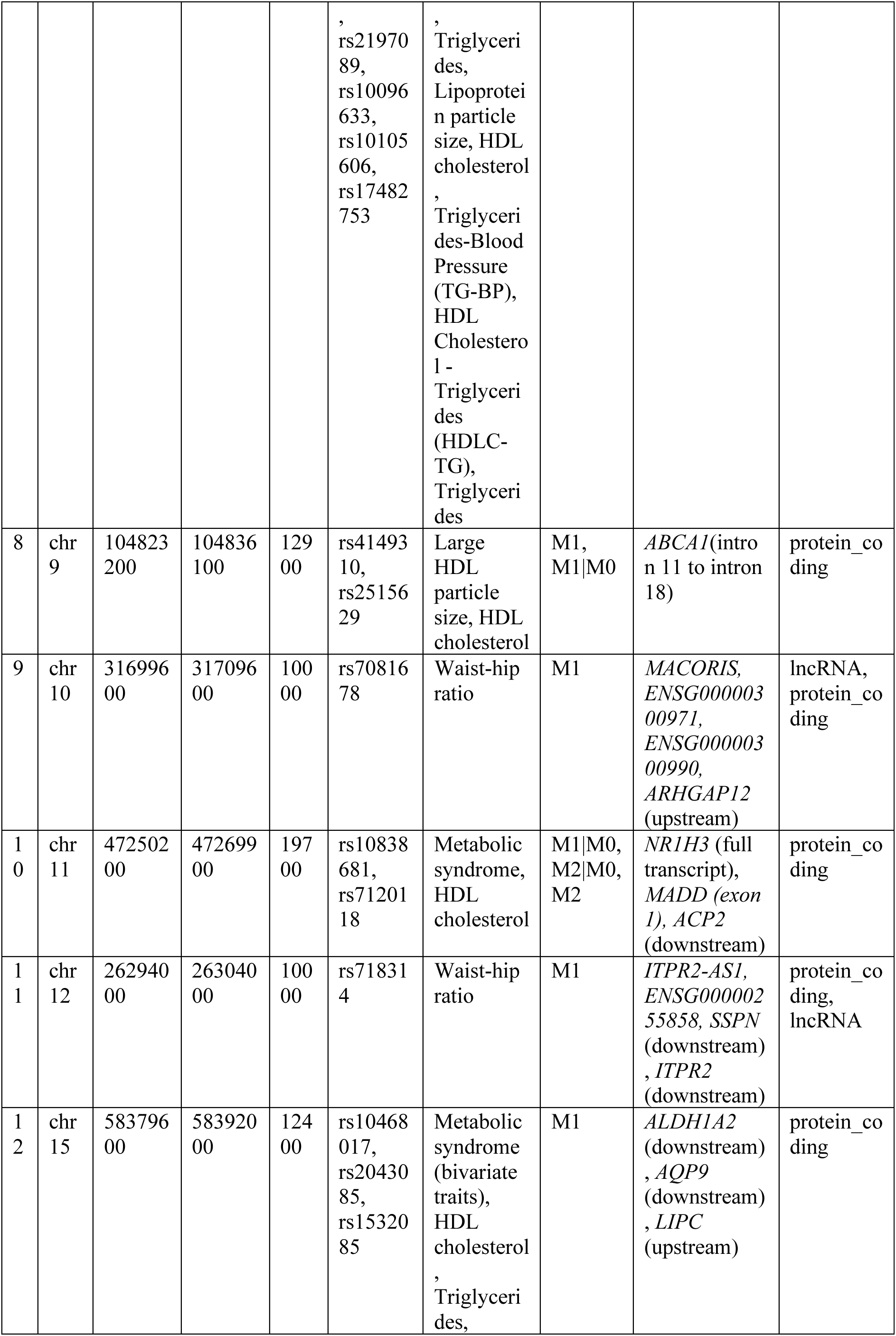
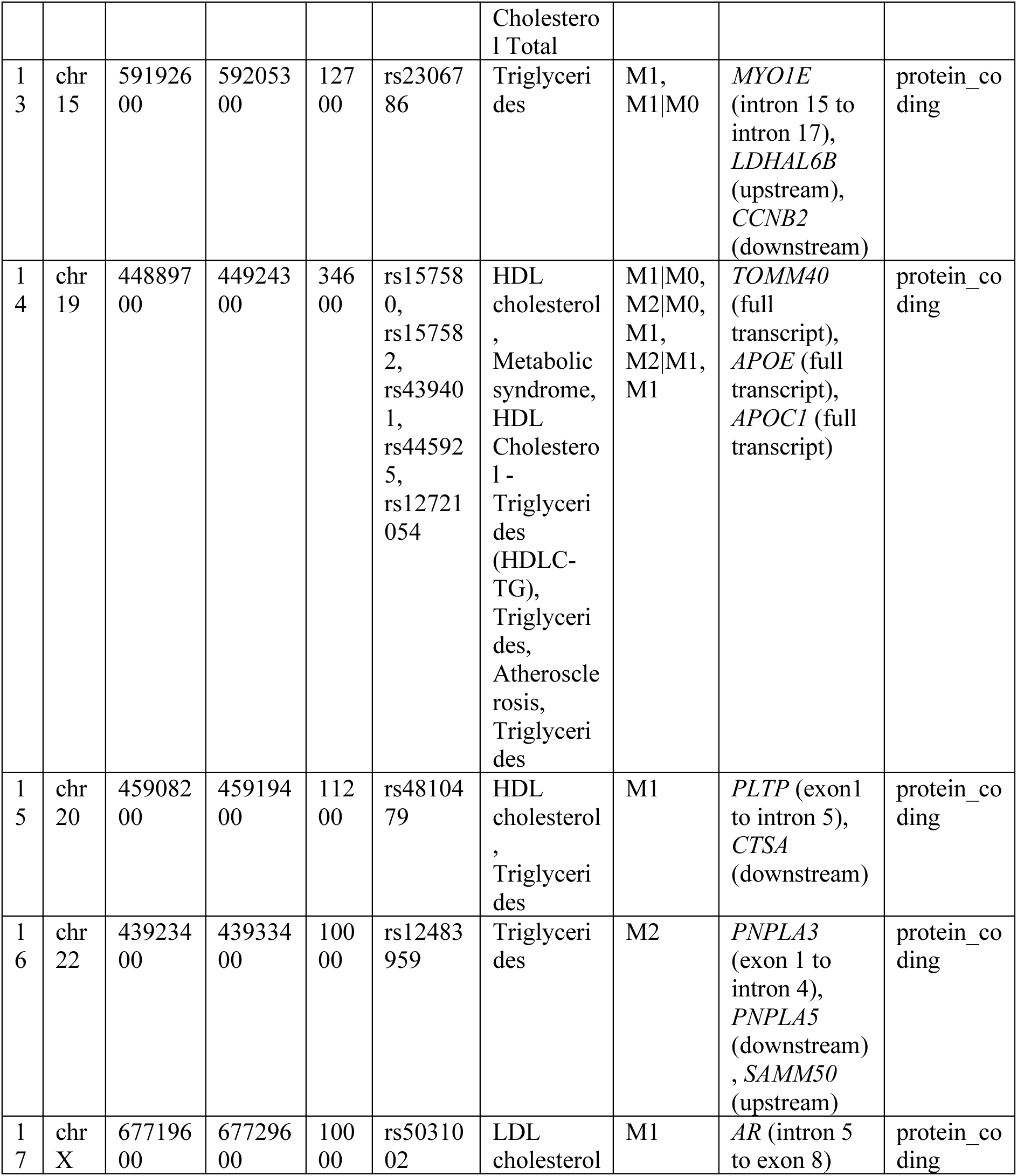
Overview of the genomic and functional features of the 17 regions enriched in macrophage enhancers and cardiovascular disease-associated SNPs. The table includes data on region coordinates, enhancer activity profiles, SNPs, GWAS traits, genes in the region. Abbreviations: Chr. = chromosome, Enh. = enhancer.

Many of these regions overlap or lie near genes with established roles in lipid metabolism, including *APOE*, *APOC1*, *LPL*, *ABCA1*, *PLTP* and *NR1H3* (also known as LXRα) (**Table 1**). These genes participate in the exogenous, endogenous and reverse cholesterol transport pathways (**Fig. S6A–C**), which have key roles in atherosclerosis formations and CVD pathogenesis [40–45]. NR1H3 acts as a transcriptional regulator of several of these genes (**Fig. S6D**) in macrophages, also crucial for the formation of atherosclerotic plaques [34,35].

In macrophages, lipoprotein lipase (LPL) hydrolyses triglycerides from circulating lipoproteins, releasing free fatty acids that can be taken up and metabolized. Beyond lipid uptake, macrophage LPL also promotes lipid accumulation and foam cell formation, contributing to atherosclerosis progression[43,46,47]. We found five non-coding CVD-related GWAS SNPs overlapping macrophage-specific genic elongating enhancers in LPL, and seven more SNPs within a newly identified enhancer region downstream of the gene (**Fig. 7A** and **Table 2**). Interestingly, the 3’ UTR rs13702 non-coding variant of LPL had previously been described as microRNA seed site that can affect RNA stability[48] but our analysis uncovered the presence of a previously unnoticed region that could also play an important regulatory role.

**Figure 7.**
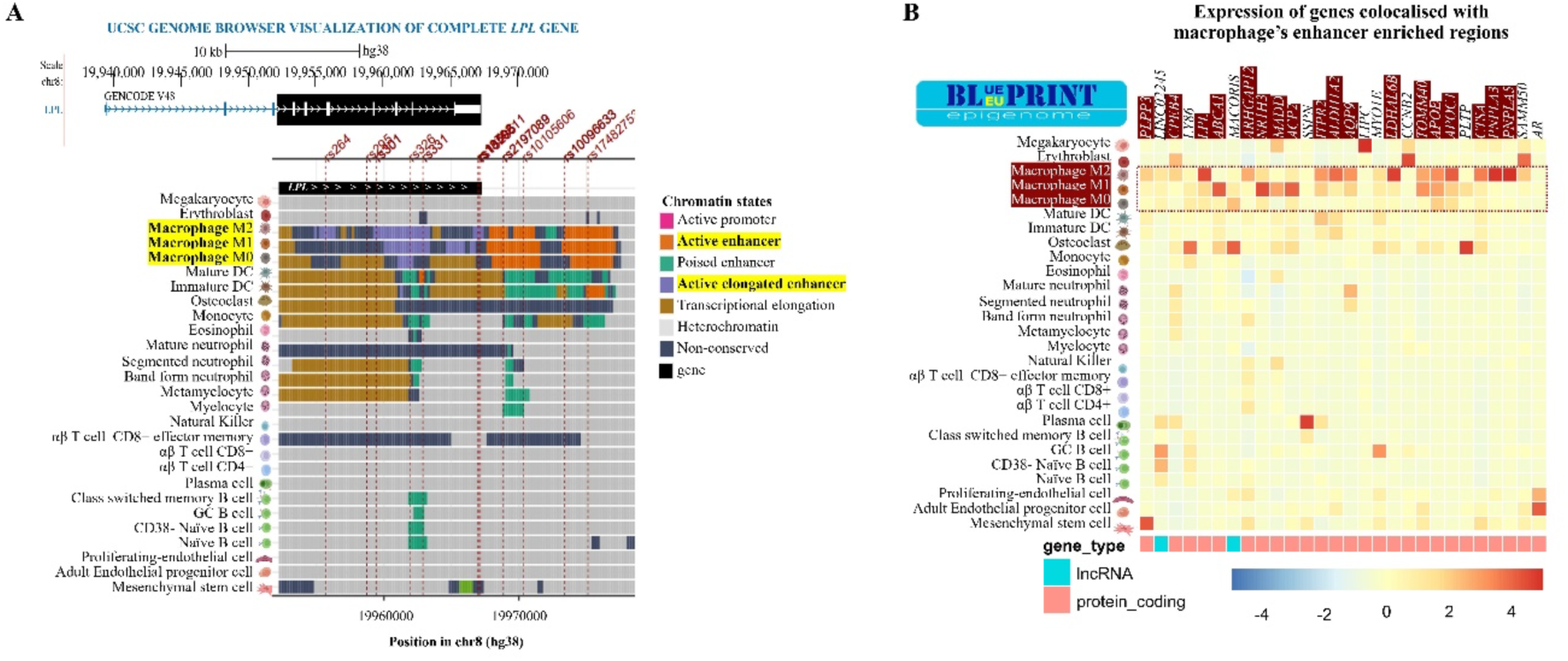
Chromatin landscape of the LPL CVD-associated enhancer locus and expression of genes overlapping CVD-associated enhancers. **(A)** Chromatin landscape of the CVD associated enhancer-enriched loci overlapping LPL gene, showing consensus chromatin states for the 31 BLUEPRINT cell types and 12 CVD-associated SNP locations (dashed vertical lines). **(B)** Heatmap with RNAseq expression data of genes overlapping or adjacent to GWAS CVD trait-associated enhancers.

**Table 2.** GWAS traits associated with non-coding SNPs in a region enriched with macrophage enhancers and overlapping LPL and NR1H3 genes.

| rsID* | Variant | GWAS trait | Gene name |
| --- | --- | --- | --- |
| rs264 | chr8_19955669_G_A | Coronary artery disease | *LPL* |
| rs295 | chr8_19958727_A_C | Metabolic syndrome | *LPL* |
| rs301 | chr8_19959423_T_C | Metabolic syndrome (bivariate traits) | *LPL* |
| rs325 | chr8_19961817_T_C | HDL cholesterol | *LPL* |
| rs326 | chr8_19961928_A_G | HDL cholesterol, Triglycerides | *LPL* |
| rs331 | chr8_19962894_G_A | Lipid metabolism phenotypes | *LPL* |
| rs13702 | chr8_19966981_T_C | HDL Cholesterol - Triglycerides (HDL-C-TG) | *LPL* |
| rs1059611 | chr8_19967052_T_C | Lipid metabolism phenotypes | *LPL* |
| rs15285 | chr8_19967156_C_T | Triglycerides-Blood Pressure (TG-BP) | *LPL* |
| rs2197089 | chr8_19968862_G_A | Metabolic syndrome (bivariate traits) | *LPL* |
| rs10105606 | chr8_19970337_C_A | Triglycerides | *LPL* |
| rs10096633 | chr8_19973410_C_T | HDL cholesterol, Metabolic traits, Triglycerides | *LPL* |
| rs17482753 | chr8_19975135_G_T | HDL cholesterol | *LPL* |
| rs2083637 | chr8_20007664_A_G | HDL cholesterol | *LPL* |
| rs1441756 | chr8_20010875_A_C | Metabolic syndrome (bivariate traits) | *LPL* |
| rs10838681 | chr11_47253513_G_A | Metabolic syndrome | *NR1H3* |
| rs7120118 | chr11_47264739_T_C | HDL cholesterol | *NR1H3* |

To assess whether these enhancer-associated genes are preferentially expressed in macrophages or more broadly across haematopoietic cell types, we analysed RNA-seq data from BLUEPRINT [49] across the 31 cell types included in our dataset (**Fig. 7B**). We identified a group of genes overlapping enhancer regions (*ABCA1, NR1H3, PNPLA3, ALDH1A2, APOE, APOC1* and *LPL*) that showed higher expression in macrophages compared to other haematopoietic lineages. We also found a second group of genes displaying high expression in macrophages but were more highly expressed in other cell types, including *CPEB4* in M2 macrophages and erythroblasts (eryth), *PLTP* in M1 macrophages and osteoclasts (osteo), and *PLPP3* in mesenchymal cells.

To extend these observations to the protein level, we examined expression patterns using the Human Protein Atlas [50]. *NR1H3*, *APOE*, *APOC1*, *LPL*, *PLTP* and *ABCA1* showed prominent gene expression across multiple macrophage populations, including circulating and tissue-resident macrophages such as Hofbauer cells, Langerhans cells and Kupffer cells **(Table S9).** These genes were also strongly expressed in metabolically relevant tissues, including blood, adipose tissue and liver. In contrast, *PNPLA3*, *PLPP3*, *CPEB4* and *ALDH1A2* were detected in macrophages but exhibited higher expression in other cell types. *PNPLA3* was predominantly expressed in liver, *PLPP3* in connective tissue and fibroblasts, *ALDH1A2* in endometrial tissues, and *CPEB4* in muscle and liver **(Table S9)**.

### Intragenic enhancers of *NR1H3* show potential distal regulatory interactions with upstream and downstream genes

The gene *NR1H3* encodes for the nuclear factor previously known as Liver X Receptor (*LXRα*). NR1H3 regulates cholesterol efflux and fatty acid metabolism in macrophages by inducing genes such as *ABCA1* and *ABCG1*, thereby maintaining lipid homeostasis [51,52]. It also suppresses pro-inflammatory gene expression through trans-repression mechanisms, linking lipid metabolism to inflammation control in macrophages[51,52]. Although this transcription factor has been studied as a regulator, we have limited information about the regulatory landscape of the locus encoding the human NR1H3 itself. Our study revealed that human macrophages and monocyte-derived DCs display a distinct chromatin landscape in the *NR1H3* locus compared to other haematological cell types, with an intra-genic active enhancer (Enh1) downstream of an active promoter region P1 (overlapping the starts of *NR1H3* and *ACP2*) upstream of an alternative active promoter (P2, **Fig. 7B**). In addition, a macrophage-specific active elongating enhancer region (Enh2) is present between P2 and the promoter (P3) of the close downstream neighbour *MADD* gene. The *NR1H3* Enh1 and Enh2 enhancers overlap with GWAS non-coding SNPs rs10838681 and rs7120118 associated with metabolic syndrome risk and HDL cholesterol levels, respectively. These CVD-associated enhancers could regulate the genes overlapping or next to them, but they could also regulate other more distal genes.

To explore the potential function of genic enhancers Enh1 and Enh2 within the gene body of *NR1H3* (**Fig. 8A**), we looked for SNPs in these regions that could be expression quantitative trait loci (eQTLs) of other genes. eQTL analysis is a method used to identify genetic variants associated with gene expression variation in the population. When applied to study enhancers, eQTL analysis can help identify genetic variants in the enhancers that can influence the expression of genes targeted by these enhancers. We refer to the genes showing significant correlations with the eQTL SNPs in enhancers as eGenes and consider them as the potential enhancer targets. Using naïve macrophages eQTL data from Alasoo et al. (2018) and Nedelec et al. (2016), reprocessed by the eQTL Catalogue[53–55], we identified 10 eGenes of SNPs within Enh1 and 3 eGenes of SNPs within Enh2 (**Fig. 8B**). Focusing on eQTL SNP-eGene interactions observed in both datasets, we see a significant interaction between Enh1 and nucleoporin *NUP160*, while for Enh2 we found interactions with *ARFGAP2* and *MYBPC3* in both studies. Interestingly, *ARFGAP2* is involved in lipid transport and vesicle trafficking.

**Figure 8.**
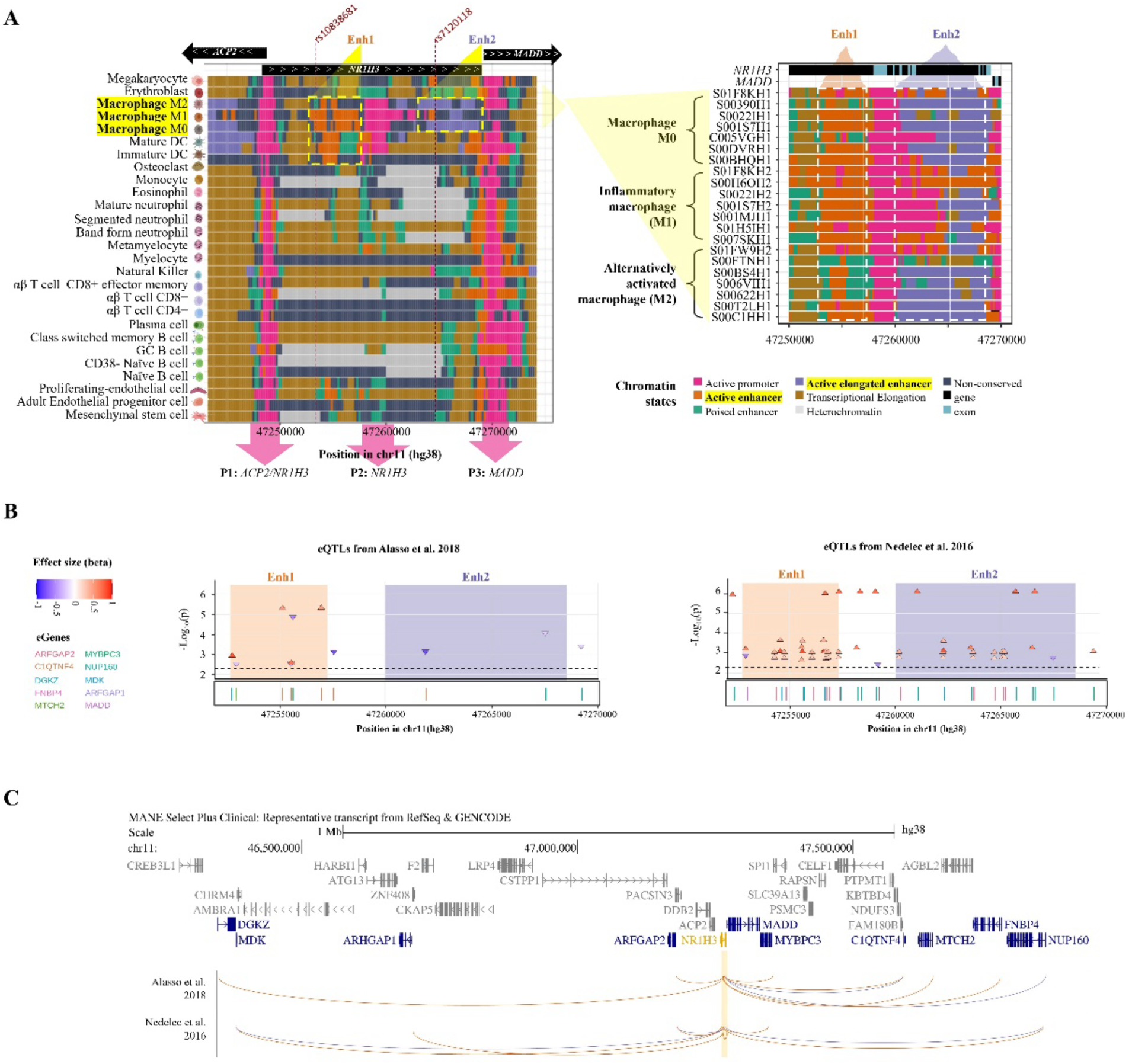
NR1H3 genic enhancers and their potential targets. **A)** Chromatin state landscape of NR1H3 locus and adjacent genes (ACP2 and MADD). The location of 2 CVD-associated SNPs (dashed vertical lines), active promoters (P1-3, pink arrows) and enhancers (Enh1, Enh2, dashed line boxes) are indicated. The left panel shows consensus chromatin states across BLUEPRINT cell types, and the right panel shows macrophage epigenomes from individual donors. **(B)** eQTL SNPs located within the genomic region described in (A), and SNP–eGene associations from Alasoo et al. (2018) and Nédélec et al. (2016) processed through the eQTL Catalogue. Points represent significant variant–gene expression associations colored according to effect size (beta). Shaded regions indicate Enh1 and Enh2. The y-axis shows statistical significance as −log10(p). **(C)** Putative enhancer–gene interactions inferred from the eQTL associations shown in (B). Curved arcs connect enhancer-associated variants to nearby eGenes across the locus.

To complement the eQTL analyses, we used the generalised Activity By Contact (gABC) predictions [56] from IHEC EpiAtlas (International Human Epigenome Consortium, in preparation, available at https://ihec-epigenomes.org/epiatlas) that identify significant correlations between H3K27ac levels in enhancers and gene expression levels in putative target genes. We observed significant co-variation of H3K27ac levels in Enh2 and changes in gene expression of *NR1H3*, *MADD*, *MAD-AS1* and *PACSIN3* genes, suggesting that this enhancer could be regulating *NR1H3* itself as well as other genes in the locus. Further experiments will be needed to validate the functionality of these enhancers and their impact on macrophages in healthy and CVD contexts, but our results strongly suggest that they might have a regulatory role in this locus.

## Discussion

Genome-wide association studies (GWAS) have uncovered thousands of genomic regions associated with complex traits [4,5], but translating these statistical associations into clear biological mechanisms remains difficult. To address this, we adopted an enhancer-centric burden approach that shifts the focus from individual variants to clusters of regulatory elements. Instead of asking whether a single variant affects gene regulation, this framework considers long enhancers or groups of enhancers linearly clustered within disease-associated loci and evaluates their activity collectively across multiple cell types. By linking non-coding variants associated with 173 complex traits to patterns of enhancer activity across combinations of haematopoietic cell types, this approach captures regulatory relationships that would be overlooked by traditional variant-centric analyses.

A key component of this analysis was the use of a consistent haematopoietic epigenomic reference dataset using data generated by the BLUEPRINT Consortium [33]. We uniformly processed 107 epigenomes from 31 primary haematopoietic cell types to derive comparable chromatin state annotations and consensus profiles. Because these data are derived from primary cells, 27 with biological replication, they offer a more physiologically relevant view of gene regulation than datasets based on immortalised cell lines. This improves the reliability of inferences about how regulatory elements function in vivo, particularly within the immune system. Although some of these cell types can be activated during the isolation processes and some cell types (including macrophages) are derived *in vitro* from primary monocytes, it represents an important resource for scientists and clinicians interested in the epigenomic regulation of the human immune system.

One of our central findings is that non-coding variant associations frequently colocalise with enhancers that are active across multiple haematopoietic cell types. This suggests that genetic risk is not confined to a single immune cellular context but is instead mediated through coordinated regulatory activity across different cell types, as it has been previously observed in the EpiMap study [9] that used the ENCODE and Roadmap datasets [15,30]. This concept is referred to as enhancer pleiotropy[57]. Such a model aligns with the multisystem nature of many complex diseases and helps explain why GWAS signals are often difficult to interpret when analysed within a single cell type or at the level of individual variants[3]. It implies that multiple regulatory elements and multiple cell types may act together to influence disease-relevant pathways.

We found that haematopoietic enhancers are not only enriched within GWAS loci associated with immune-related traits but also for metabolic, cardiovascular, neurological, and cancer-related phenotypes. This broader pattern reflects the role of the immune system as a central regulator of physiological processes throughout the body. It suggests that immune regulatory programs may contribute more widely to the genetic architecture of complex traits than previously recognised, extending beyond classical immune or inflammatory diseases.

This highlights a limitation of previous studies [9–16] that, although successful in linking GWAS variants to regulatory annotations, have largely relied on reference datasets such as those generated by the ENCODE Project and the NIH Roadmap Epigenomics Project [15,30]. Although hugely valuable, these resources have limited representation of immune cell types. By incorporating a more comprehensive set of haematopoietic epigenomes from the BLUEPRINT project [33], this study extends existing variant-to-regulatory annotation frameworks, enabling a more detailed and context-specific interpretation of non-coding genetic variation in complex traits.

However, several limitations of our study should be considered. Enhancer activity and enhancer–gene links are inferred from chromatin state data and integrative models and therefore require experimental validation. The activity profiles are likely incomplete, as regulatory elements active in unprofiled cell types or conditions are not captured. Functional validation in primary human cells remains challenging and requires coordinated clinical access to relevant samples, molecular expertise in enhancer perturbation, and robust phenotypic readouts[60,61]. These capabilities are rarely co-located within a single research setting, and the separation between computational and experimental domains, together with the substantial time and resource requirements of validation, continues to limit the translation of large-scale genomic analyses into mechanistic insight.

To illustrate how this framework can help bridge the gap between genetic association and disease mechanisms, we focused on cardiovascular disease (CVD) and related traits. Our results showed that a substantial fraction of CVD-associated loci is linked to macrophage enhancer activity and overlaps genes with well-established roles in lipid metabolism, including *APOE, APOC1, LPL, ABCA1, PLTP* and *NR1H3*. These genes participate in pathways regulating lipid uptake, transport and cholesterol homeostasis and are expressed in both macrophages and metabolically relevant tissues [40–45]. Given the central role of macrophages in atherosclerosis [34,35,58], the identification of CVD-associated enhancers linked to lipid metabolism genes places these regulatory elements within pathways directly relevant to disease pathogenesis. Our analysis further delineates specific enhancer regions within these loci that may regulate these genes, providing candidate regulatory elements for functional investigation and supporting a model in which non-coding variants may influence disease risk through macrophage lipid-handling programmes.

At the locus level, analysis of *NR1H3* illustrates the complexity of regulatory architecture underlying GWAS signals. We identify intragenic enhancers associated with GWAS variants and link these elements to candidate target genes using macrophage eQTL [53–55] data and a generalised activity-by-contact model [56,59]. These results suggest that non-coding variation at this locus may influence both local gene regulation and distal transcriptional programmes. Although these approaches are particularly sensitive to distal interactions, they do not exclude proximal regulatory effects, and regulation of *NR1H3* itself and neighbouring genes is likely to represent an important component of enhancer function at this locus. Future analyses of the transcription factor landscape of the enhancers regions we have identified will provide more detailed mechanistic interpretation. Together, these findings highlight how multiple variants within enhancer regions may act in combination to influence disease-relevant pathways within specific cellular contexts.

In summary, our results support a model in which the effects of non-coding genetic variation are best understood within a multi-cell-type regulatory framework. Considering enhancer activity as coordinated units across cellular contexts provides a richer and more coherent interpretation of GWAS signals and supports a shift from variant-centric to regulatory architecture-based models of genetic risk. Although experimental validation is beyond the scope of this study, we provide this resource as a foundation for further work by groups with the appropriate experimental and clinical capabilities to investigate these regulatory mechanisms.

## Material and methods

### Sources for ChIP-Seq data acquisition

ChIP-seq data used in this study were generated by the BLUEPRINT Consortium [33]. Sequence alignments to the human reference genome (hg38/GRCh38) were performed by the BLUEPRINT Data Coordination Centre (DCC). Aligned data were downloaded from the European Genome-phenome Archive (accession EGAS00001000326) [62]. A total of 642 ChIP-seq experiments and 107 input controls were retrieved as BAM files and converted to BED format for downstream analysis. Briefly, alignments were performed using BWA 0.7.7 [63], Picard v2.8.1 (https://broadinstitute.github.io/picard/), and samtools v1.3.1 [64]. Duplicated reads were removed and quality of the data was assessed using PhantomPeakQualTools [65].

The dataset includes the whole-genome profiling of six histone marks (H3K4me1, H3K4me3, H3K27me3, H3K36me3, H3K9me3, H3K27ac), as well as corresponding inputs/controls, for 107 human samples from 31 different blood-derived cell types belonging to the main hematopoietic lineages: myeloid (15 cell types) and lymphoid (12 cell types). These datasets also include endothelial (2 cell types) and bone marrow mesenchymal cells (2 cell types) **(Fig 1. A-D).** Sample metadata, such as sex, tissue, biomaterial type (primary or primary derived), cell type, and EpiRR accessions, are included in **Table S1.**

### Learning the haematopoietic chromatin state maps

Chromatin state maps published by independent subgroups within the BLUEPRINT Consortium [66–73] are not directly comparable due to differences in sample subsets, computational methods, and genome assemblies used across studies. Here we addressed this by generating comparable chromatin state maps for BLUEPRINT samples. We trained and applied a 12-state chromatin model to a collection of 642 ChIP-seq alignments to generate 107 epigenomic maps, including maps for cell types not previously reported.

Chromatin state modelling was performed using ChromHMM (v1.10) [74], based on multivariate Hidden Markov Models. The model was trained on six core histone modifications (H3K4me1, H3K4me3, H3K27me3, H3K36me3, H3K9me3 and H3K27ac) together with matched input controls, default ChromHMM parameters were used.

The number of states was set to 12 to balance interpretability and model complexity. Other ChromHMM models based on similar datasets, including the 18- and 15-state models from the Roadmap Consortium [15] and the 11-state model by Carrillo *et al.* [38], have been described previously. The 11-state model captures key biologically interpretable states (including active promoter, bivalent promoter, enhancer, elongation, and heterochromatin/low signal) represented in the larger models but does not include genic enhancer states. The 12-state model used here retains these core states while additionally capturing genic enhancer states.

The trained model was applied to each sample to estimate the posterior probability of each chromatin state across the genome, and genomic 200 bp bins were assigned to the most probable state to generate genome-wide chromatin state maps.

### State labels, interpretation, and mnemonics

To assign biologically meaningful mnemonics to the states of the 12-state model, we followed the approach of Carrillo *et al.* (2017). Emission probabilities were cross-correlated with those from the 11-state model [38], and Roadmap models (18- and 15-state) [15] (see **Tables S2, S3** and **Fig. S1A–B**). States with correlation coefficients >0.75 were considered equivalent (**Fig. S1C**).

The 12-state model captures most states identified in larger models, with the exception of Flanking TSS Downstream, Weak Enhancer, Bivalent/Poised TSS, and Bivalent Enhancer.

Additionally, following the approach of Carrillo *et al.* (2017) [38], the 12-state model was collapsed into seven interpretable chromatin states based on literature and genomic annotation enrichments (including gene structure and CpG islands). The resulting states were: Polycomb (states 1–2), Heterochromatin (HetChrom; states 3–4), Transcriptional Elongation (TransElo; states 5–6), Elongating Active Enhancer (AleloE; state 7), Poised Enhancer (PoiseE; state 8), Conventional Active Enhancer (AconE; states 9–10), and Active Promoter (Aprom; states 11–12). This approach to collapsing chromatin states for interpretability has been widely adopted in previous studies [38,66,67,71].

### Cell type consensus epigenomes

To compare chromatin states across epigenomes, we constructed a matrix of ChromHMM state annotations with genomic bins as rows and samples as columns. Chromosome-specific annotations were concatenated by stacking bins in genomic order across chromosomes, and for each bin the chromatin state annotation was recorded across the 107 samples. The resulting matrix comprised 14,374,996 bins × 107 samples.

Consensus chromatin states were defined by states by condensing ChromHMM annotations across biological replicates for each cell type, following previously described approaches [67,71]. Only cell types with at least two biological replicates were included. For each genomic bin, a consensus state was assigned if the same annotation was observed in at least 75% of biological replicates of the same cell type; otherwise, the bin was classified as non-conserved within that cell type. This threshold was selected to balance robustness of state assignment with variability in replicate number across cell types.

### Enhancer activity profiles

Active enhancer (AE) bins (200 bp) were defined as genomic bins annotated as elongating or conventional active enhancers (AleloE or AconE states) in at least two of the 107 epigenomes, resulting in 1,526,184 AE bins. This definition captures enhancer activity across individual epigenomes rather than relying on consensus annotations.

For each AE bin, an activity profile was defined as the set of cell types in which the bin was annotated as a conserved active enhancer in the corresponding consensus epigenomes. AE bins were then grouped by shared activity profiles, irrespective of genomic location. This resulted in 107,562 distinct enhancer groups, which varied in size, with most groups (75%) comprising a small number of bins.

### GWAS enrichment analysis

To investigate the functional relevance of non-coding trait-associated variants, we integrated data from the GWAS catalogue [4] with active enhancer (AE) bins identified in this study. SNPs annotated as protein-altering variants (missense, stop-gain, frameshift) were excluded, retaining variants located in non-coding or regulatory regions("intergenic", "", "ncRNA", "cds-synon", "intron", "UTR-5", "UTR-3", "splice-3", "nearGene-", "nearGene-5", or "splice-5"). Analyses were restricted to studies with more than 2,000 individuals and traits associated with at least six SNPs, resulting in 11,060 SNPs across 518 traits (985 PubMed IDs).

Trait-associated regions were defined as 10 kb windows centred on each GWAS SNP (±5 kb). Enrichment analyses focused on AE bins belonging to the 100 most frequent activity profiles across the 31 BLUEPRINT cell types.

Overlaps between AE bins and trait-associated regions were computed, and enrichment of specific enhancer activity profiles was assessed using Fisher’s exact test, comparing bins with a given activity profile to all other AE bins as background. P-values were adjusted for multiple testing using the Bonferroni method, with a significance threshold of adjusted *P* < 1 × 10⁻⁴.

### Gene and protein expression datasets

RNA-seq data generated by the BLUEPRINT project were used to calculate average gene expression across replicates for the 31 cell types. These data were obtained from the final BLUEPRINT data release (July 2016), as no subsequent releases were made by the consortium. The datasets are available from: https://ftp.ebi.ac.uk/pub/databases/blueprint/data/homo_sapiens/GRCh38/, and accompanying data descriptions can be found in in https://ftp.ebi.ac.uk/pub/databases/blueprint/. Gene expression values were also obtained from obtained from the Human Protein Atlas [50].

### eQTL and enhancer–gene association analyses

Expression quantitative trait loci (eQTL) analyses were used to identify associations between genetic variants located within enhancer regions and gene expression. SNPs overlapping enhancer regions were mapped to associated genes (eGenes) using naïve macrophage eQTL datasets from Alasoo *et al.* (2018) and Nédélec *et al.* (2016), as reprocessed by the eQTL Catalogue [53–55]. Only SNP–gene associations replicated across both datasets were retained.

To complement eQTL-based associations, enhancer–gene links were further evaluated using generalised Activity-by-Contact (gABC) predictions obtained from the (https://ihec-epigenomes.org/epiatlas/)[75]. These predictions integrate enhancer activity (H3K27ac signal) and gene expression to infer putative regulatory interactions between enhancers and target genes) [56,59]. We used the gABC_StackedChromHMM_InteractionsSampleMat.txt dataset and applied an enhancer activity threshold cutoff of >0.02. The specific version of the file used in this study is available at https://doi.org/10.6084/m9.figshare.32231184.

### Software and code availability

All analyses were performed using R version 3.6.3 (2020-02-29) [76]. Scripts used in this study are available at https://github.com/juearcilaga/Unmasking-disease-risk-in-haematopoietic-enhancers.

## Data Availability

The datasets generated during this study are available in the Figshare repository (https://doi.org/10.6084/m9.figshare.25601301). These include chromatin state maps for 107 BLUEPRINT donors, consensus chromatin state profiles for 31 haematopoietic cell types, active enhancer annotations, GWAS enrichment results, and SNP-to-enhancer mapping datasets. The repository also contains all reference input datasets used during the analysis, including processed gene annotations, transcript expression data, and associated BLUEPRINT, RNA-seq, and ChIP-seq metadata files.

The code used for data processing and analysis is publicly available at GitHub repository https://github.com/juearcilaga/Unmasking-disease-risk-in-haematopoietic-enhancers.

## Supplementary Figures

**Figure S1.**
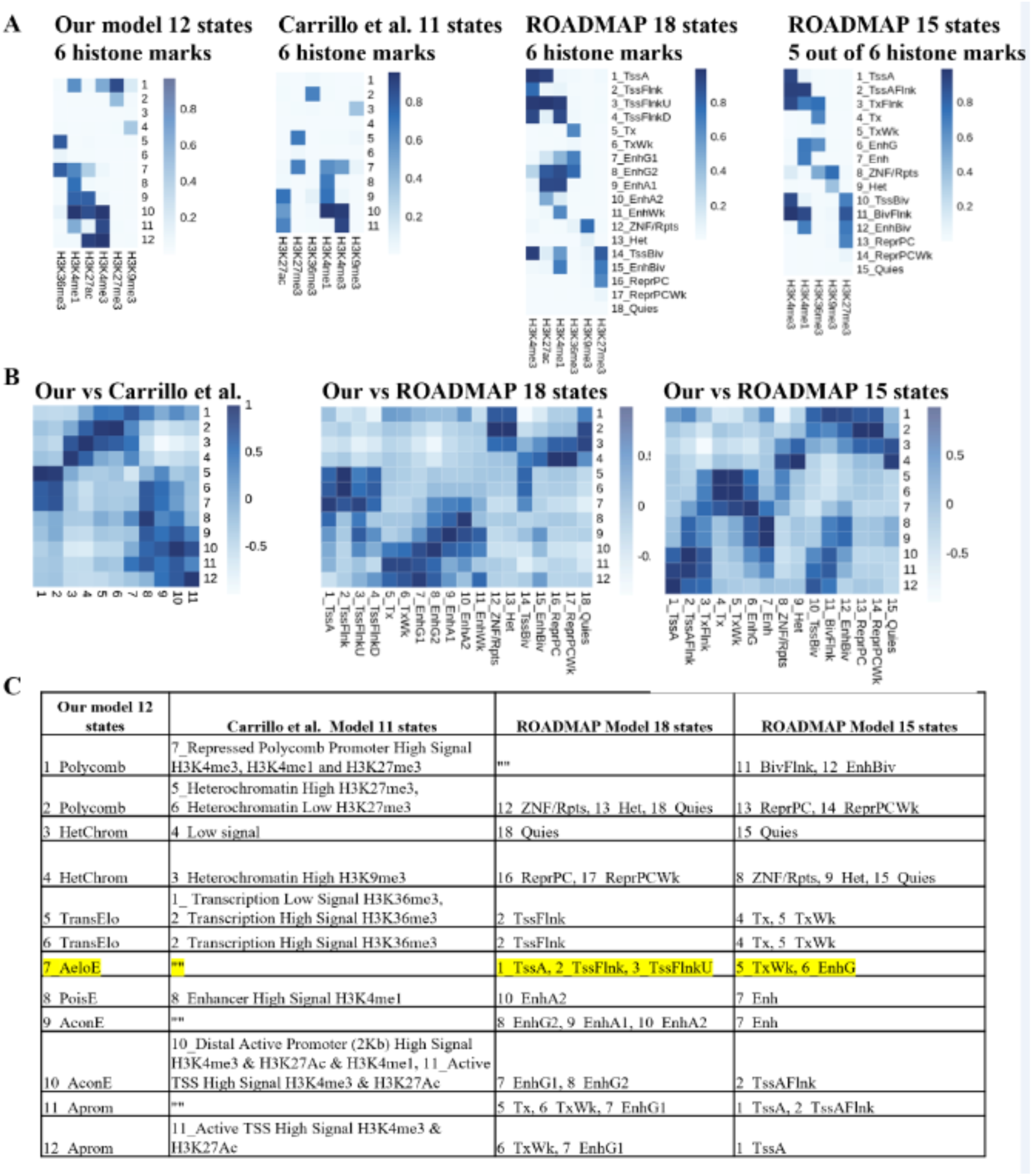
Correlation between the emission probabilities of our model (12 states) and those of corresponding states from previously published models. (A) Shows the emission probabilities of our 12-states model, the 11-states model from Carrillo et al. (2017), and those from the 18 and 15-state models from NIHR Roadmap. **(B)** Heatmap displaying the coefficient values from the Pearson Correlation between our model and the 11-states model by Carrillo et al. (2017), as well as the 18-states and 15-states models by the NIH Epigenomic Roadmap. (C) Best match (correlation coefficient > 0.75) between each state in our 12 states-model and the states from Carrillo et al. (2017) and NIHR Roadmap (2015) models. The figure displays the abbreviations of the chromatin state names used in the original papers; see Tables S2 and S3 for full names.

**Figure S2.**
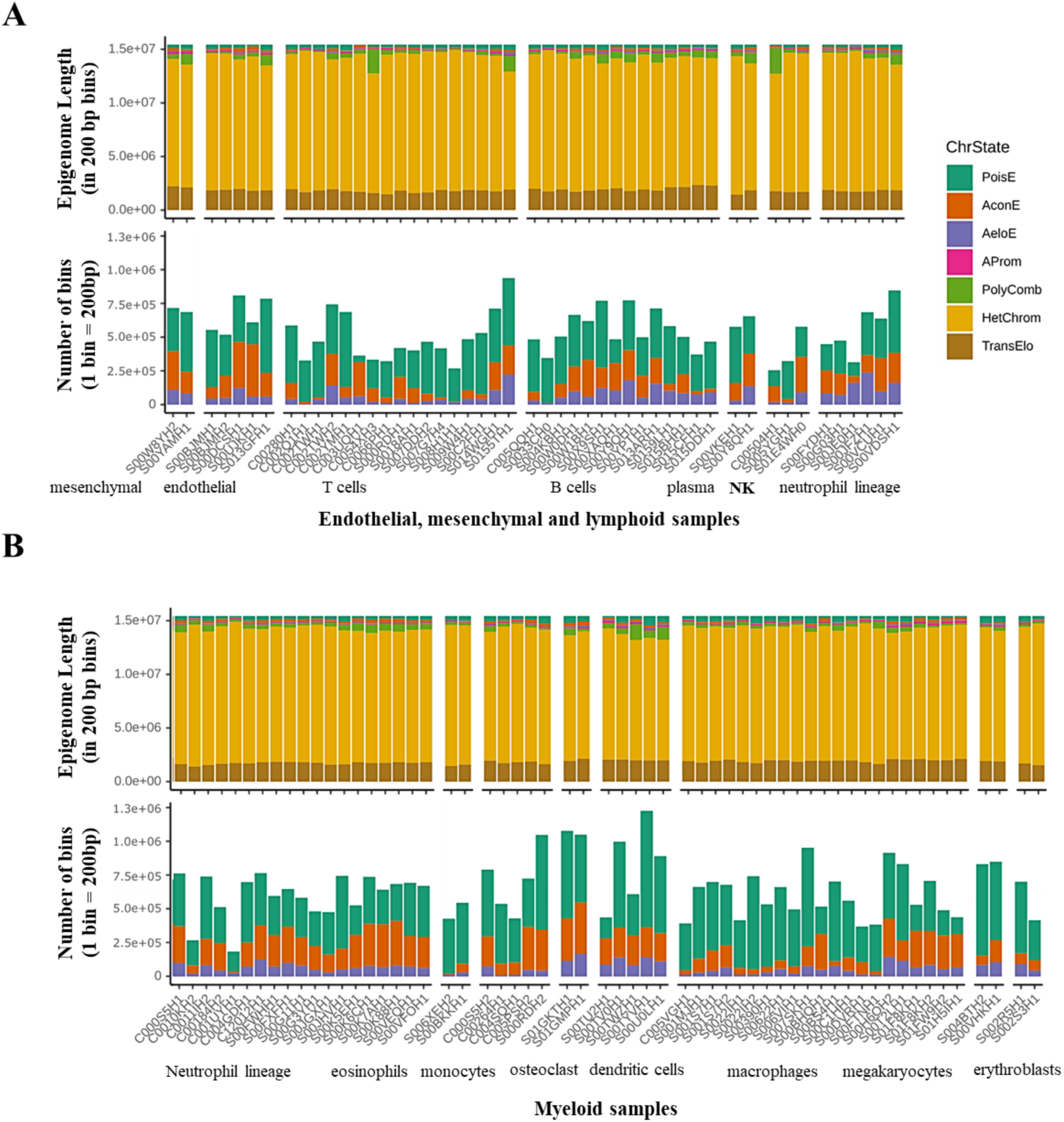
Epigenomic profile of haematopoietic cell types from the BLUEPRINT Consortium. **(A)** Endothelial, mesenchymal and lymphoid cell types **(B)** Myeloid cell types. For each of the 107 haematopoietic epigenomes generated in this study, the bar plots represent: The proportion of the epigenome length covered by each chromatin state (Top panel). The number of bins (1 bin = 200bp) annotated as each of the three types of enhancers (Bottom panel).

**Figure S3.**
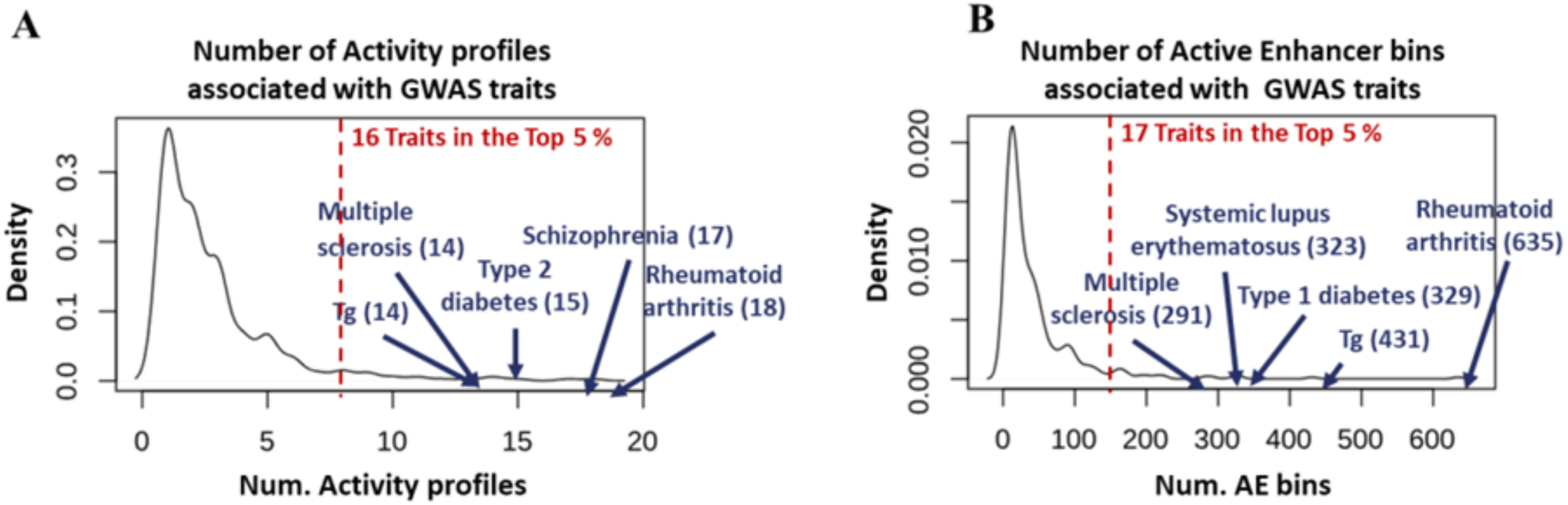
Interplay between genetic susceptibility, enhancer dynamics, and cellular contexts in complex diseases. The figure illustrates that a single trait can be associated with numerous enhancers, each exhibiting unique activity profiles across various cell types.Density plots depict the distribution of different features linked to GWAS traits: **(A)** the number of activity profiles, **(B)** active enhancer bins. The red dotted line indicates the density point until 95% of the traits are covered. The number of traits within the upper 5% is highlighted in bold red font, emphasising a subset of 5 traits within the most extreme values, denoted in bold blue font.

***Figure S4 goes as a separate PDF file.***

***Figure S4. Cell types associated with each group of complex traits.*** *The heatmap illustrates the various cell types participating in the activity profiles of AE associated with each group of GWAS traits including **(A)** Immune-response-related, **(B)** Cardio and Metabolic and Lipid, **(C)** Haemmatological **(D)** Neuro **(E)** Substance use, **(F)** Eye, **(G)** Cancer, **(H)** Anthropometric, **(I)** Biochemical, **(J)** Other*

***Figure S5 goes as a separate PDF file.***

***Figure S5. Chromatin landscape around the cardiovascular disease-associated loci enriched in macrophage enhancers.***

**Figure S6.**
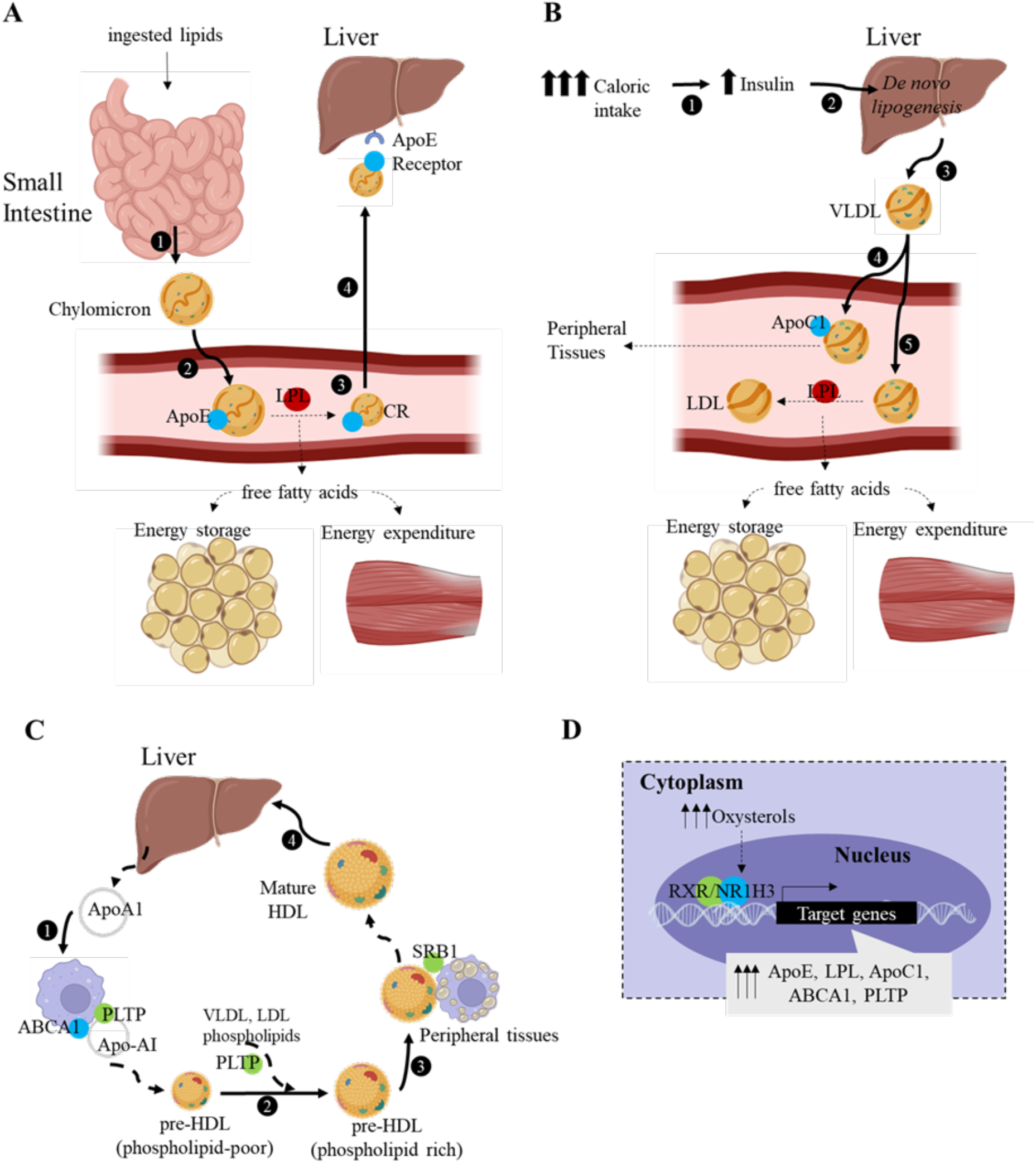
Mapping macrophage enhancers target genes to specific steps in lipid metabolism. This figure shows a simplified version of the lipid metabolism pathways, highlighting the role of the proteins encoded by the genes potentially regulated by macrophage enhancers associated with CVD risk. **(A)** Exogenous Pathway: In the intestine, ingested lipids are packed into chylomicrons by enterocytes (Step 1). As these particles reach the bloodstream, they acquire ApoE (Step 2). Upon reaching target tissues, LPL hydrolyses their content, facilitating tissue uptake for storage or energy production (Step 3). Post lipolysis, chylomicrons become chylomicron remnants (CR) (Step 4). These remnants are recognised by hepatocytes via an ApoE receptor and cleared from the bloodstream (Step 5). **(B)** Endogenous Pathway: Triggered by excessive caloric intake, insulin concentration in the blood increases; this pathway begins with insulin signalling the liver to start lipogenesis (Step 1). Liver-synthesized lipids are packaged into VLDL (Step 2), which, in circulation, can undergo lipolysis in target tissues by LPL (Step 3) or acquire APOC1, inhibiting LPL and allowing VLDL to travel further to peripheral tissues (Step 4). **(C).** Reverse Cholesterol (RCT) Pathway: ApoAI synthesised by the liver acquires phospholipids from macrophages via ABCA1 and PLTP (Step 1). Remodelled particles become preHDL, acquiring more phospholipids from other lipoproteins in circulation and increasing their size (Step 2). Phospholipids from PreHDL particles sequester cholesterol effluxed from macrophages via SRB1 (Step 3) and, after collecting cholesterol, become mature HDL particles that are cleared from the bloodstream by the liver (Step 4). **D.** NHR1H3, also known as LXR alpha, has a master regulatory role in orchestrating these pathways, this transcription factor regulates the transcription of ABCA1, APOE, LPL, APOC1, PLTP and other several genes that are key players in lipid metabolism.

