## Supplementary figures and images for "An enhancer-centric approach applied to human immune system epigenomes revealed the association of macrophage enhancers with cardiovascular disease"

### Figure S4

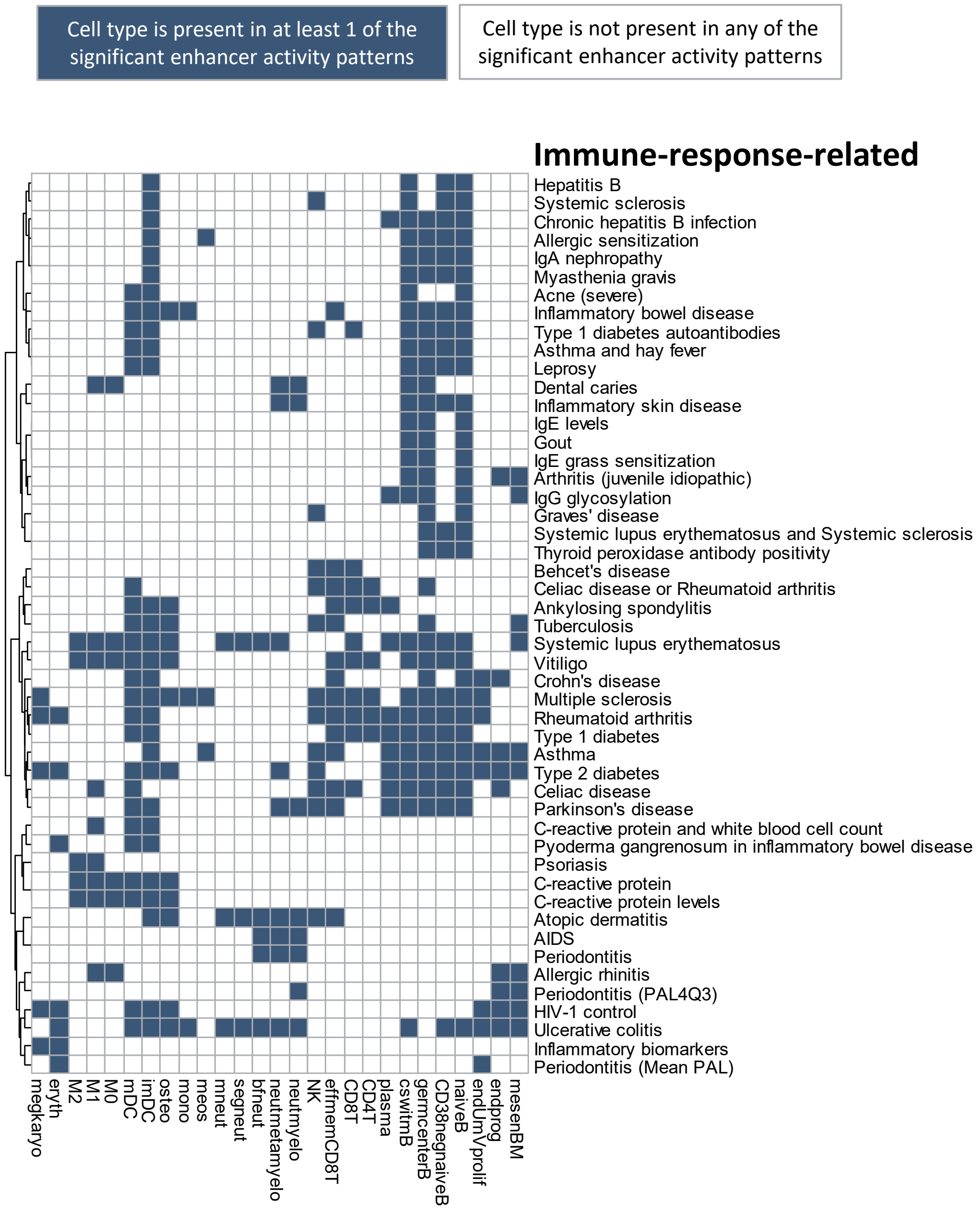

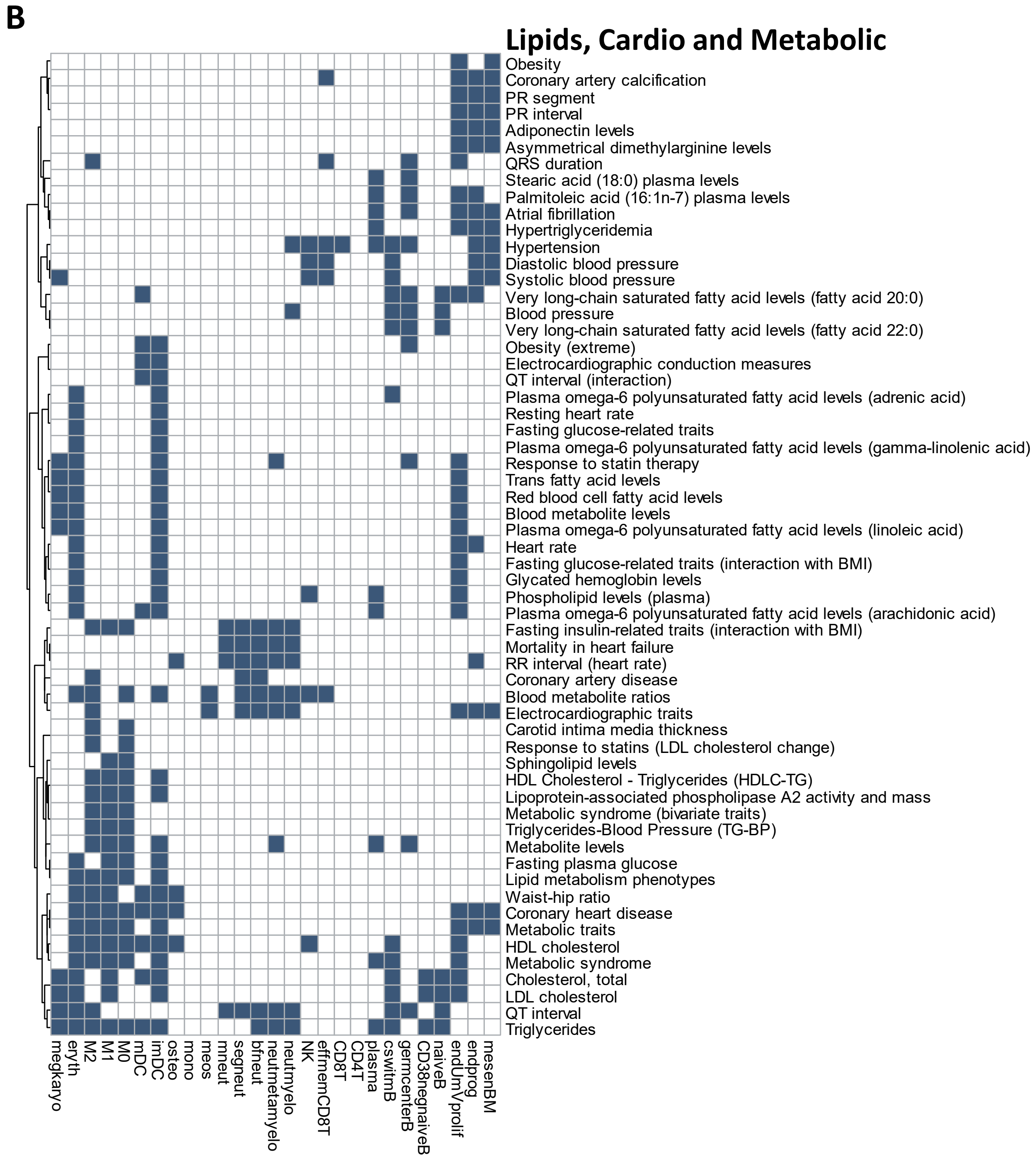

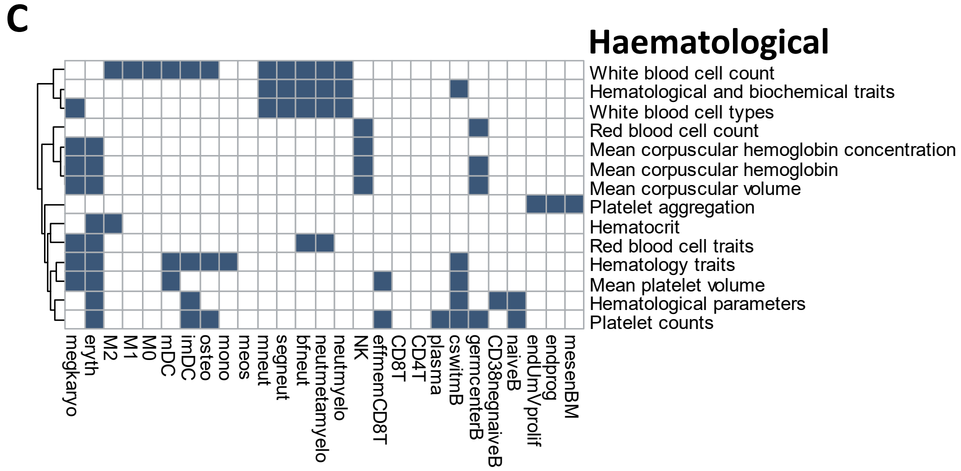

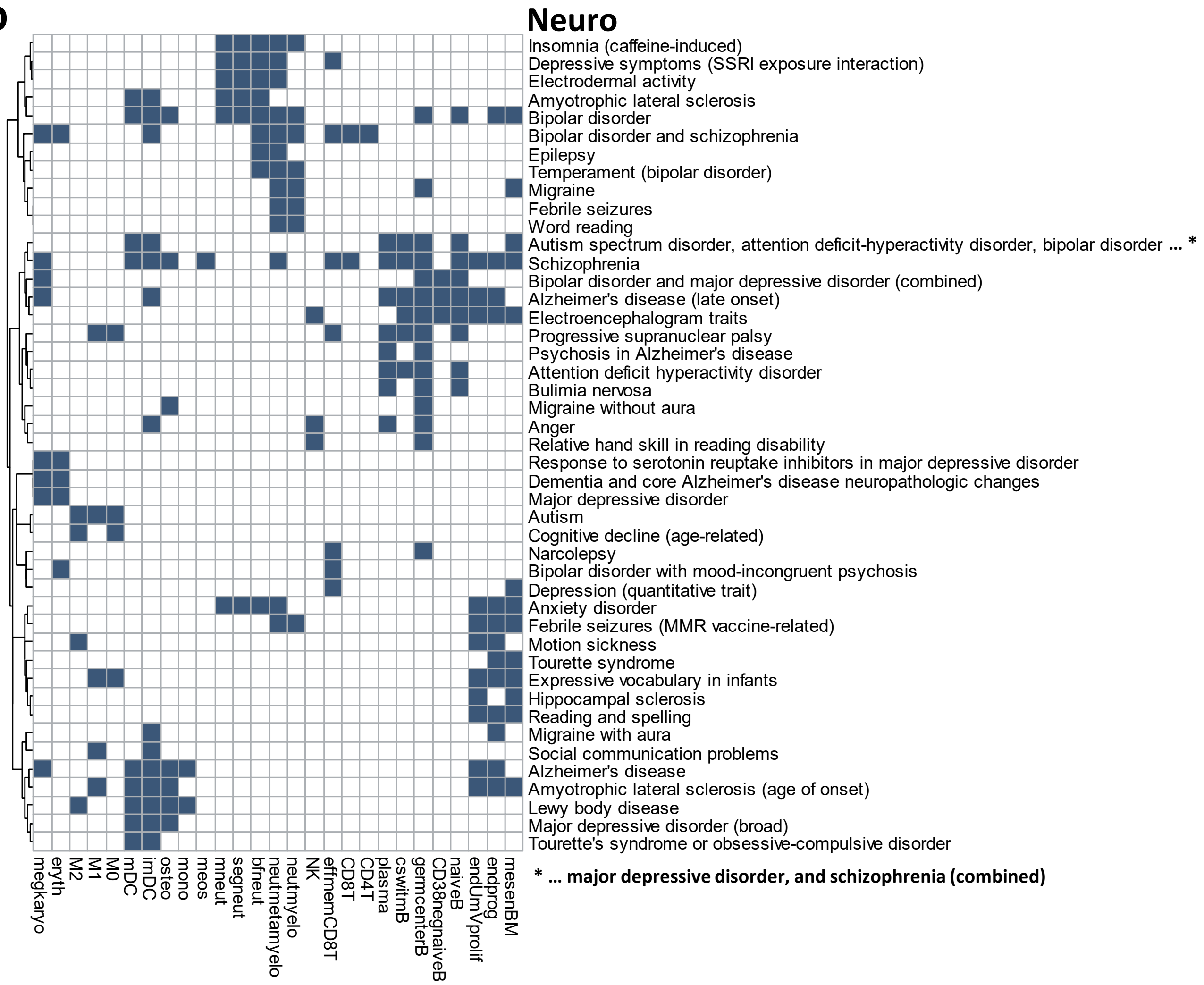

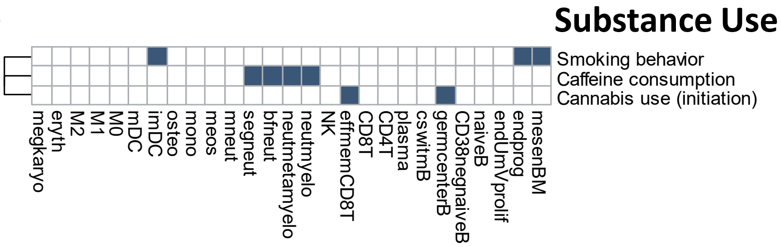

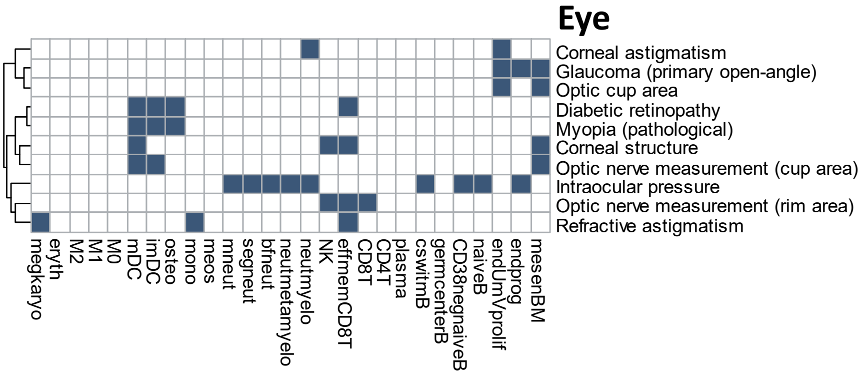

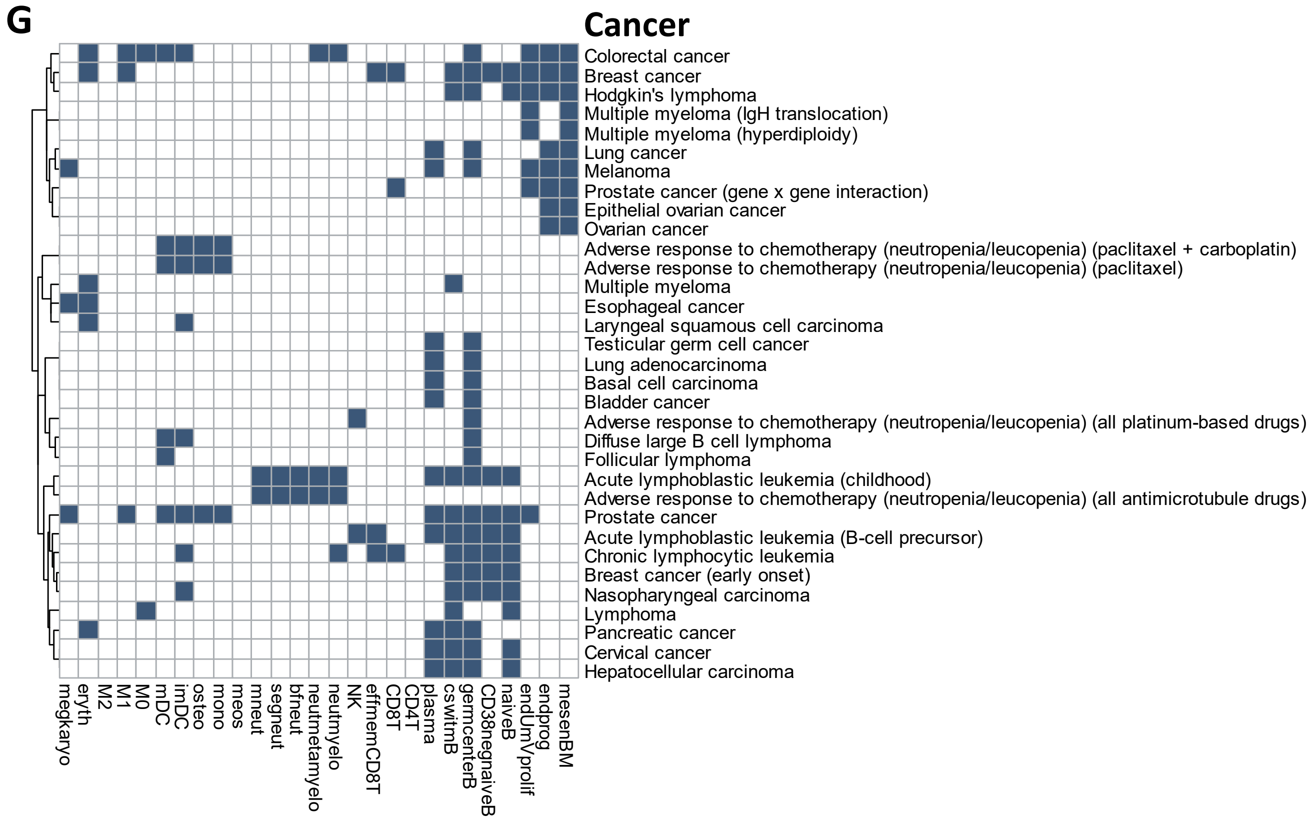

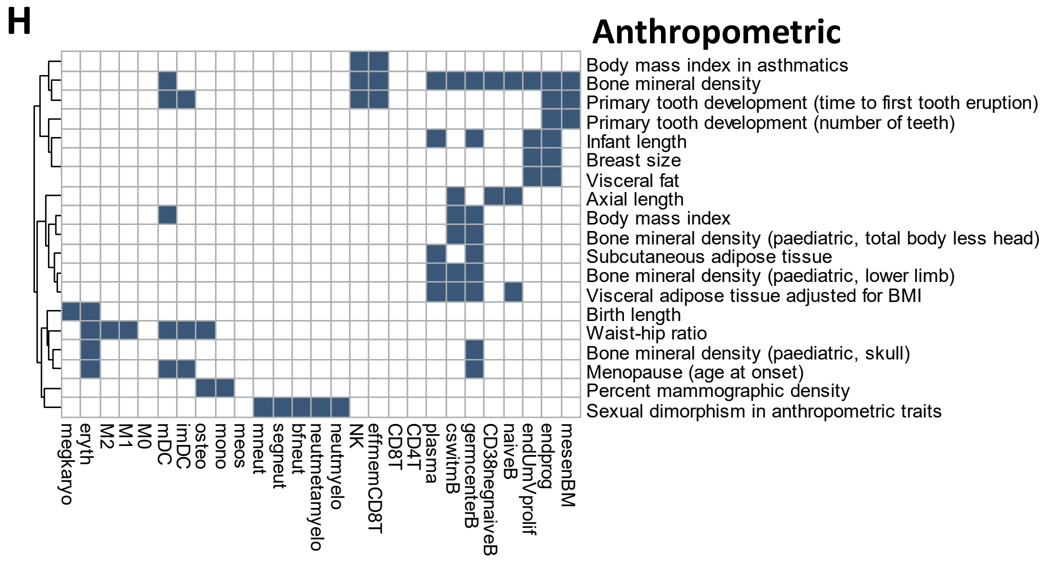

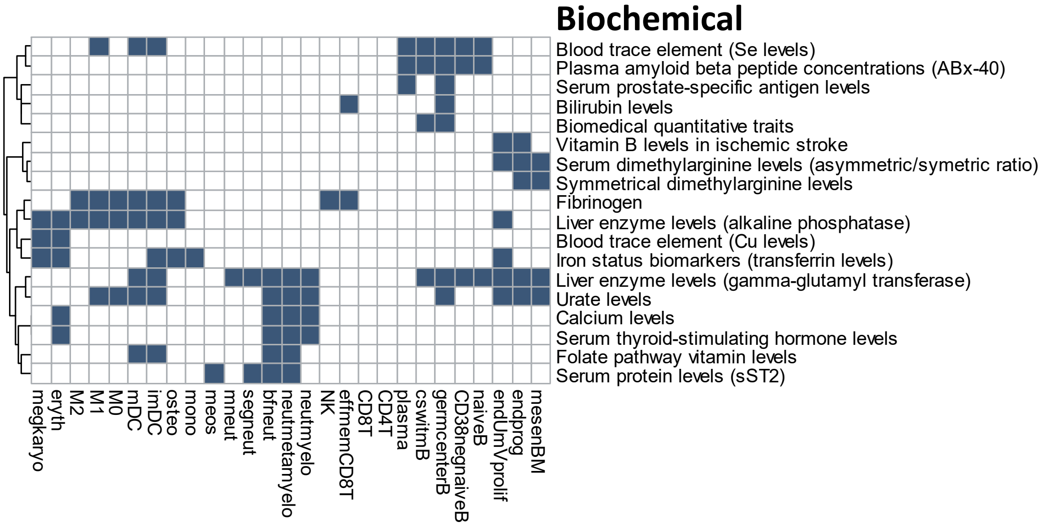

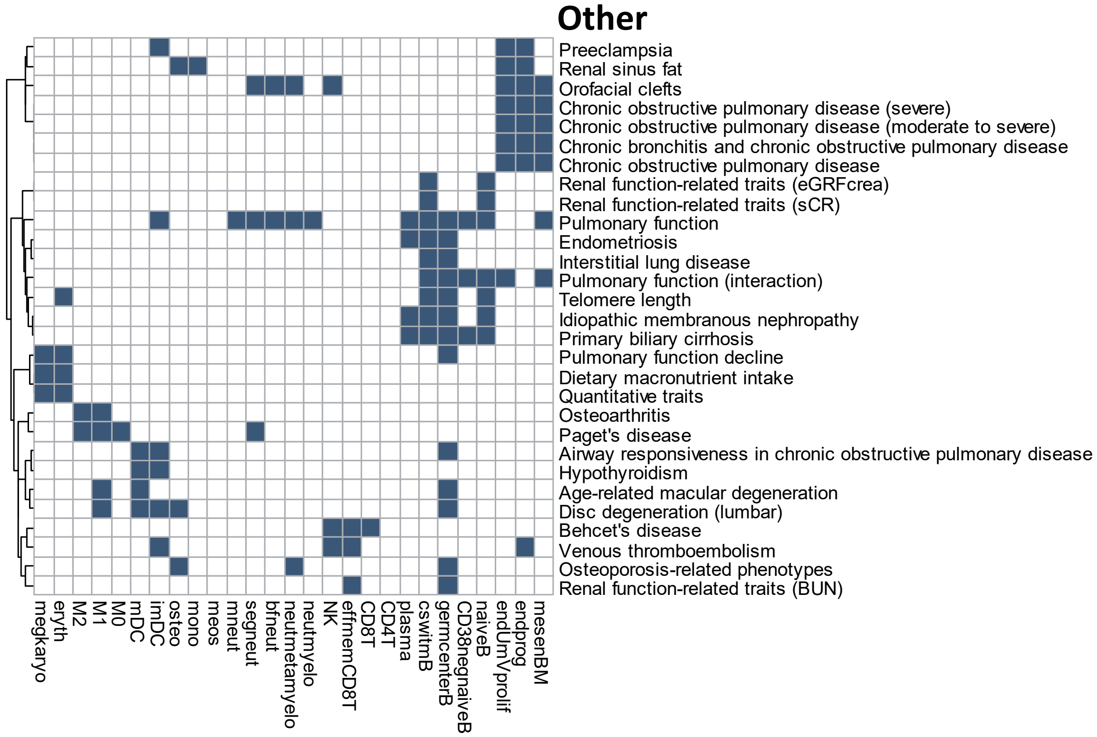

### Figure S5

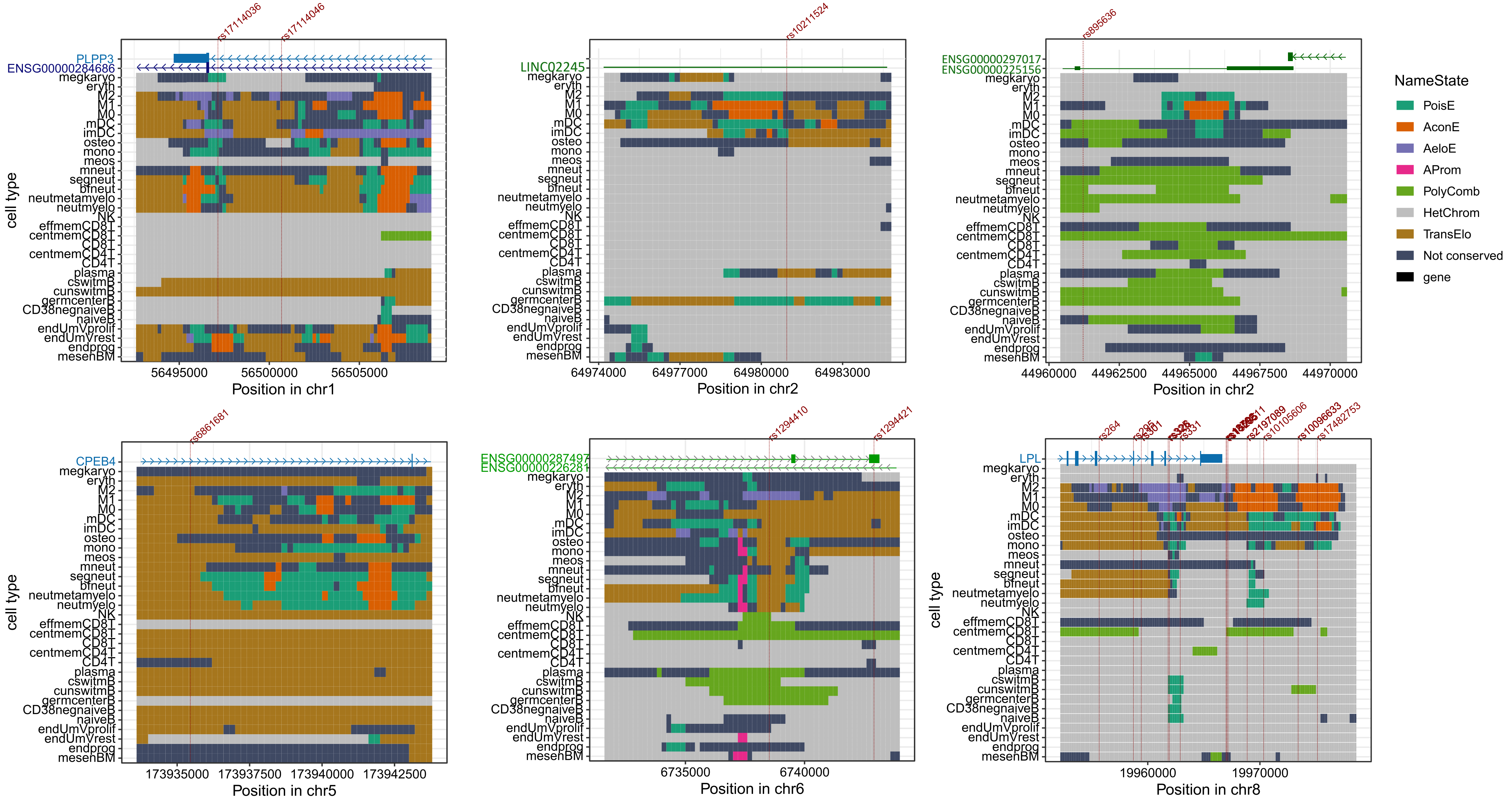

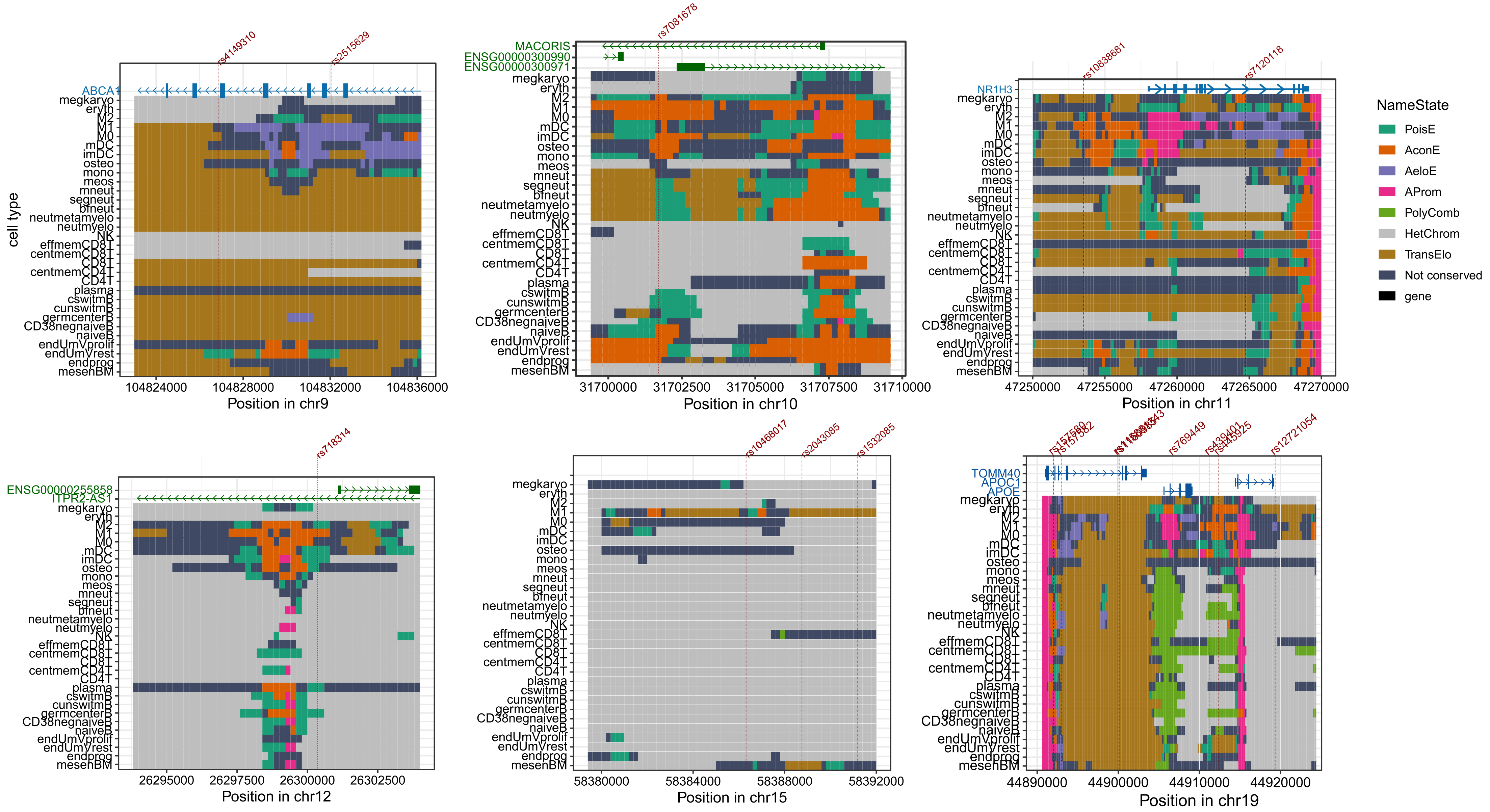

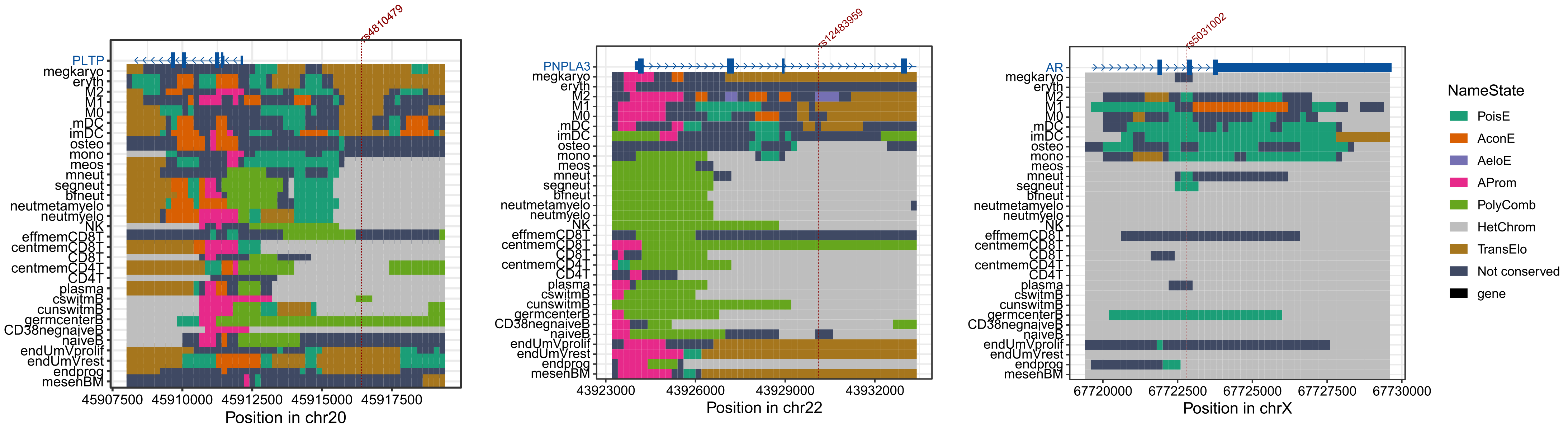
