## Supplementary material for "An enhancer-centric approach applied to human immune system epigenomes revealed the association of macrophage enhancers with cardiovascular disease": Tables S1, S2, S8 and S9

Table S1. Sample metadata.

| **BIOMATERIAL TYPE** | **SAMPLE NAME** | **CELL TYPE** | **TISSUE TYPE** | **DONOR SEX** | **DONOR ID** | **EPIRR** |
| --- | --- | --- | --- | --- | --- | --- |
| Primary Cell Culture | S00622H1 | alternatively activated macrophage | venous blood | Male | S00622 | IHECRE00000129.3 |
| Primary Cell Culture | S006VIH1 | alternatively activated macrophage | venous blood | Male | S006VI | IHECRE00000059.3 |
| Primary Cell Culture | S00BS4H1 | alternatively activated macrophage | venous blood | Female | S00BS4 | IHECRE00000013.3 |
| Primary Cell Culture | S00C1HH1 | alternatively activated macrophage | cord blood | Female | S00C1H | IHECRE00000055.3 |
| Primary Cell Culture | S00FTNH1 | alternatively activated macrophage | venous blood | Female | S00FTN | IHECRE00000071.3 |
| Primary Cell Culture | S00T2LH1 | alternatively activated macrophage | cord blood | Male | S00T2L | IHECRE00000269.2 |
| Primary Cell Culture | S01FW9H2 | alternatively activated macrophage | venous blood | Male | S01FW9 | IHECRE00001391.1 |
| Primary Cell Culture | S004BTH2 | CD34-negative, CD41-positive, CD42-positive megakaryocyte cell | cord blood | Female | S004BT | IHECRE00000105.3 |
| Primary Cell Culture | S00VHKH1 | CD34-negative, CD41-positive, CD42-positive megakaryocyte cell | cord blood | Male | S00VHK | IHECRE00000257.2 |
| Primary Cell Culture | S00BJMH1 | endothelial cell of umbilical vein (proliferating) | cord blood | Male | S00BJM | IHECRE00000184.3 |
| Primary Cell Culture | S00DCSH1 | endothelial cell of umbilical vein (proliferating) | cord blood | Female | S00DCS | IHECRE00000126.3 |
| Primary Cell Culture | S00BJMH2 | endothelial cell of umbilical vein (resting) | cord blood | Male | S00BJM | IHECRE00000099.3 |
| Primary Cell Culture | S002R5H1 | erythroblast | cord blood | Male | S002R5 | IHECRE00000193.3 |
| Primary Cell Culture | S002S3H1 | erythroblast | cord blood | Female | S002S3 | IHECRE00000112.3 |
| Primary Cell Culture | S00TU2H1 | immature conventional dendritic cell | venous blood | Male | B270 | IHECRE00001548.1 |
| Primary Cell Culture | S00TV0H1 | immature conventional dendritic cell | venous blood | Male | B271 | IHECRE00001457.1 |
| Primary Cell Culture | S00TWZH1 | immature conventional dendritic cell | venous blood | Male | B272 | IHECRE00001387.1 |
| Primary Cell Culture | S001MJH1 | inflammatory macrophage | venous blood | Male | S001MJ | IHECRE00000174.3 |
| Primary Cell Culture | S001S7H2 | inflammatory macrophage | venous blood | Female | S001S7 | IHECRE00000043.3 |
| Primary Cell Culture | S0022IH2 | inflammatory macrophage | venous blood | Female | S0022I | IHECRE00000161.3 |
| Primary Cell Culture | S007SKH1 | inflammatory macrophage | cord blood | Male | S007SK | IHECRE00000195.3 |
| Primary Cell Culture | S00H6OH2 | inflammatory macrophage | venous blood | Male | S00H6O | IHECRE00000318.2 |
| Primary Cell Culture | S01F8KH2 | inflammatory macrophage | venous blood | Male | S01F8K | IHECRE00001293.1 |
| Primary Cell Culture | S01H5IH1 | inflammatory macrophage | cord blood | Female | S01H5I | IHECRE00001289.1 |
| Primary Cell Culture | S001S7H1 | macrophage | venous blood | Female | S001S7 | IHECRE00000084.3 |
| Primary Cell Culture | S0022IH1 | macrophage | venous blood | Female | S0022I | IHECRE00000177.3 |
| Primary Cell Culture | S00390H1 | macrophage | venous blood | Male | S00390 | IHECRE00000008.3 |
| Primary Cell Culture | S00BHQH1 | macrophage | cord blood | Female | S00BHQ | IHECRE00000121.3 |
| Primary Cell Culture | S00DVRH1 | macrophage | cord blood | Male | S00DVR | IHECRE00000060.3 |
| Primary Cell Culture | S01F8KH1 | macrophage | venous blood | Male | S01F8K | IHECRE00001398.1 |
| Primary Cell Culture | C005VGH1 | macrophage | venous blood | Male | C005VG |  |
| Primary Cell Culture | S00TYVH1 | mature conventional dendritic cell | venous blood | Male | B271 | IHECRE00001546.1 |
| Primary Cell Culture | S00U0LH1 | mature conventional dendritic cell | venous blood | Male | B272 | IHECRE00001490.1 |
| Primary Cell Culture | S01GKTH1 | osteoclast | venous blood | Male | BC2_0 | IHECRE00001249.1 |
| Primary Cell Culture | S01GMPH1 | osteoclast | venous blood | Male | BC2_10 | IHECRE00001471.1 |
| Primary Cell | S00UJKH1 | adult endothelial progenitor cell | venous blood | Female | S00UJK | IHECRE00000303.2 |
| Primary Cell | S013GFH1 | adult endothelial progenitor cell | venous blood | Female | S013GF | IHECRE00001510.1 |
| Primary Cell | S00VEQH1 | band form neutrophil | bone marrow | Female | BM060814 | IHECRE00000285.2 |
| Primary Cell | S00G11H1 | band form neutrophil | bone marrow | Male | BM220513 |  |
| Primary Cell | S00JGXH1 | band form neutrophil | bone marrow | Male | BM030613 |  |
| Primary Cell | C000S5H2 | CD14-positive, CD16-negative classical monocyte | venous blood | Male | C000S5 | IHECRE00000027.3 |
| Primary Cell | C00264H1 | CD14-positive, CD16-negative classical monocyte | cord blood | Male | C00264 | IHECRE00000135.3 |
| Primary Cell | C004SQH1 | CD14-positive, CD16-negative classical monocyte | venous blood | Female | C004SQ | IHECRE00000101.3 |
| Primary Cell | C005PSH2 | CD14-positive, CD16-negative classical monocyte | cord blood | Female | C005PS | IHECRE00000155.3 |
| Primary Cell | S000RDH2 | CD14-positive, CD16-negative classical monocyte | cord blood | Male | S000RD | IHECRE00000048.3 |
| Primary Cell | S004KBH1 | CD38-negative naive B cell | venous blood | Male | S004KB | IHECRE00000125.3 |
| Primary Cell | C005QQH1 | CD38-negative naive B cell | cord blood | Female | C005QQ |  |
| Primary Cell | S0033CH0 | CD38-negative naive B cell | venous blood | Male | C003JB,C003RW,C003N3,C003QY | |
| Primary Cell | C00280H1 | CD4-positive, alpha-beta T cell | cord blood | Female | C00280 | IHECRE00001251.1 |
| Primary Cell | C002Q1H1 | CD4-positive, alpha-beta T cell | venous blood | Male | C002Q1 | IHECRE00000009.3 |
| Primary Cell | C002TWH1 | CD4-positive, alpha-beta T cell | venous blood | Male | C002TW | IHECRE00000075.3 |
| Primary Cell | S000RDH1 | CD4-positive, alpha-beta T cell | cord blood | Male | S000RD | IHECRE00000140.3 |
| Primary Cell | S0018AH1 | CD4-positive, alpha-beta T cell | cord blood | Female | S0018A | IHECRE00000191.3 |
| Primary Cell | S008H1H1 | CD4-positive, alpha-beta T cell | venous blood | Male | S008H1 | IHECRE00000160.3 |
| Primary Cell | S009W4H1 | CD4-positive, alpha-beta T cell | venous blood | Female | S009W4 | IHECRE00000194.3 |
| Primary Cell | S007DDH2 | CD4-positive, alpha-beta T cell | venous blood | Female | S007DD |  |
| Primary Cell | S007G7H4 | CD4-positive, alpha-beta T cell | venous blood | Male | S007G7 |  |
| Primary Cell | C002YMH1 | CD8-positive, alpha-beta T cell | cord blood | Female | C002YM | IHECRE00000035.3 |
| Primary Cell | C0066PH1 | CD8-positive, alpha-beta T cell | cord blood | Female | C0066P | IHECRE00000076.3 |
| Primary Cell | S00C2FH1 | CD8-positive, alpha-beta T cell | cord blood | Male | S00C2F | IHECRE00000022.3 |
| Primary Cell | S014WGH1 | CD8-positive, alpha-beta T cell | venous blood | Female | S014WG |  |
| Primary Cell | C002TWH2 | central memory CD4-positive, alpha-beta T cell | venous blood | Male | C002TW | IHECRE00000102.3 |
| Primary Cell | S0155TH1 | central memory CD8-positive, alpha-beta T cell | venous blood | Male | S0155T |  |
| Primary Cell | S00YPTH1 | class switched memory B cell | venous blood | Male | S00YPT | IHECRE00001540.1 |
| Primary Cell | S015BHH1 | class switched memory B cell | venous blood | Male | csMBC pool 2 | IHECRE00001255.1 |
| Primary Cell | S015CFH1 | class switched memory B cell | venous blood | Female | csMBC pool 8 |  |
| Primary Cell | S005YGH1 | cytotoxic CD56-dim natural killer cell | cord blood | Male | S005YG | IHECRE00000049.3 |
| Primary Cell | C00504H1 | cytotoxic CD56-dim natural killer cell | venous blood | Female | C00504 |  |
| Primary Cell | S01E4WH0 | cytotoxic CD56-dim natural killer cell | cord blood | Female | S01DWH,S01DWH,S01DWH,S01DWH,S01DWH | |
| Primary Cell | C003UQH1 | effector memory CD8-positive, alpha-beta T cell | venous blood | Male | C003UQ | IHECRE00000010.3 |
| Primary Cell | C0054XH3 | effector memory CD8-positive, alpha-beta T cell | venous blood | Female | C0054X | IHECRE00000017.3 |
| Primary Cell | S00Y9OH1 | germinal center B cell | tonsil | Female | T14_10 | IHECRE00000332.2 |
| Primary Cell | S013ARH1 | germinal center B cell | tonsil | Male | T14_11 | IHECRE00001375.1 |
| Primary Cell | S00W0DH1 | germinal center B cell | tonsil | Female | T14_5 |  |
| Primary Cell | S00BKKH1 | mature eosinophil | venous blood | Female | S00BKK | IHECRE00000114.3 |
| Primary Cell | S006XEH2 | mature eosinophil | venous blood | Male | S006XE |  |
| Primary Cell | C000S5H1 | mature neutrophil | venous blood | Male | C000S5 | IHECRE00000109.3 |
| Primary Cell | C0010KH2 | mature neutrophil | venous blood | Female | C0010K | IHECRE00000004.3 |
| Primary Cell | C0011IH2 | mature neutrophil | venous blood | Female | C0011I | IHECRE00000159.3 |
| Primary Cell | C00184H2 | mature neutrophil | cord blood | Male | C00184 | IHECRE00000095.3 |
| Primary Cell | C001UYH1 | mature neutrophil | venous blood | Male | C001UY | IHECRE00000094.3 |
| Primary Cell | C004GDH1 | mature neutrophil | cord blood | Female | C004GD | IHECRE00000124.3 |
| Primary Cell | C12012H1 | mature neutrophil | venous blood | Male | C12012 | IHECRE00000178.3 |
| Primary Cell | S00FWHH1 | mature neutrophil | venous blood | Male | PB130513 |  |
| Primary Cell | S00FXFH1 | mature neutrophil | venous blood | Male | PB130513 |  |
| Primary Cell | S00K5EH1 | mature neutrophil | venous blood | Male | PB100713 |  |
| Primary Cell | S00K6CH1 | mature neutrophil | venous blood | Male | PB100713 |  |
| Primary Cell | S00K7AH1 | mature neutrophil | venous blood | Male | PB270313 |  |
| Primary Cell | S00K88H1 | mature neutrophil | venous blood | Male | PB270313 |  |
| Primary Cell | S00W8YH2 | mesenchymal stem cell of the bone marrow | venous blood | Unknown | S00W8Y | IHECRE00001335.1 |
| Primary Cell | S00YAMH1 | mesenchymal stem cell of the bone marrow | venous blood | Unknown | S00YAM | IHECRE00000250.2 |
| Primary Cell | S00X9SH1 | naive B cell | venous blood | Male | NC14_42 | IHECRE00000280.2 |
| Primary Cell | S00XAQH1 | naive B cell | venous blood | Male | NC14_47 | IHECRE00000258.2 |
| Primary Cell | S00W1BH1 | naive B cell | venous blood | Male | NC14_5 |  |
| Primary Cell | S0159LH1 | naive B cell | venous blood | Female | B15_50 |  |
| Primary Cell | S00G03H1 | neutrophilic metamyelocyte | bone marrow | Male | BM220513 | IHECRE00000262.2 |
| Primary Cell | S00VDSH1 | neutrophilic metamyelocyte | bone marrow | Female | BM060814 | IHECRE00000248.2 |
| Primary Cell | S00JFZH1 | neutrophilic metamyelocyte | bone marrow | Male | BM030613 |  |
| Primary Cell | S00VCUH1 | neutrophilic myelocyte | bone marrow | Female | BM060814 | IHECRE00000330.2 |
| Primary Cell | S00FYDH1 | neutrophilic myelocyte | bone marrow | Male | BM220513 |  |
| Primary Cell | S00JE0H1 | neutrophilic myelocyte | bone marrow | Male | BM030613 |  |
| Primary Cell | S00Y8QH1 | plasma cell | tonsil | Female | T14_10 | IHECRE00000347.2 |
| Primary Cell | S00VKEH1 | plasma cell | tonsil | Female | T14_5 |  |
| Primary Cell | S00VFOH1 | segmented neutrophil of bone marrow | bone marrow | Female | BM060814 | IHECRE00000317.2 |
| Primary Cell | S00G3YH1 | segmented neutrophil of bone marrow | bone marrow | Male | BM220513 |  |
| Primary Cell | S00JHVH1 | segmented neutrophil of bone marrow | bone marrow | Male | BM030613 |  |
| Primary Cell | S015DDH1 | unswitched memory B cell | venous blood | Male | Pool_9 |  |

* Rows highlighted in yellow indicate samples that have not been included in the EPIRR.

Table S2. Correlation of emission probabilities between Carrillo et al. (2017) model and ours.

| **state from Carrillo et.al (2017)** | **state mnemonic (as in original paper)** | **state label description (as in original paper)** | **best match* (correlation coefficient)** | **best match* (state from our model)** |
| --- | --- | --- | --- | --- |
| 1 | Transcription | Transcription Low Signal H3K36me3 | 0.74 | 12 |
| 2 | Transcription | Transcription High Signal H3K36me3 | 0.63 | 9 |
| 3 | Heterochromatin | Heterochromatin High Signal H3K9me3 | 1 | 4 |
| 4 | Heterochromatin | Low signal | 0.95 | 4 |
| 5 | Heterochromatin | Heterochromatin High Signal H3K27me3 | 1 | 8 |
| 6 | Heterochromatin | Heterochromatin Low Signal H3K27me3 | 1 | 8 |
| 7 | Repressed Promoter | Repressed Polycomb Promoter High Signal H3K4me3, H3K4me1 and H3K27me3 | 0.89 | 1 |
| 8 | Enhancer | Enhancer High Signal H3K4me1 | 0.85 | 11 |
| 9 | Enhancer | Active Enhancer High Signal H3K4me1 & H3K27Ac | 0.62 | 5 |
| 10 | Active Promoter | Distal Active Promoter (2Kb) High Signal H3K4me3 & H3K27Ac & H3K4me1 | 0.54 | 2 |
| 11 | Active Promoter | Active TSS High Signal H3K4me3 & H3K27Ac | 0.76 | 2 |

*Top value for the Pearson correlation coefficient between each state (11 states) in Carrillo et al. (2027) model and our model (12 states). States with correlation coefficients below 0.75 are highlighted in red.

Table S3. Correlation of emission probabilities between NIHR ROADMAP (2015) model and ours.

| **state from ROADMAP (2015)** | **state mnemonic (as in original paper)** | **state label description (as in original paper)** | **best match* (correlation coefficient)** | **best match* (state from our model)** |
| --- | --- | --- | --- | --- |
| 1 | TssA | Active TSS | 0.94 | 7 |
| 2 | TssFlnk | Flanking TSS | 1 | 6 |
| 3 | TssFlnkU | Flanking TSS Upstream | 0.89 | 7 |
| 4 | TssFlnkD | Flanking TSS Downstream | 0.69 | 6 |
| 5 | Tx | Strong transcription | 0.87 | 11 |
| 6 | TxWk | Weak transcription | 0.86 | 11 |
| 7 | EnhG1 | Genic enhancer1 | 0.94 | 12 |
| 8 | EnhG2 | Genic enhancer2 | 0.97 | 10 |
| 9 | EnhA1 | Active Enhancer 1 | 1 | 9 |
| 10 | EnhA2 | Active Enhancer 2 | 0.97 | 8 |
| 11 | EnhWk | Weak Enhancer | 0.68 | 9 |
| 12 | ZNF/Rpts | ZNF genes & repeats | 0.99 | 2 |
| 13 | Het | Heterochromatin | 1 | 2 |
| 14 | TssBiv | Bivalent/Poised TSS | 0.63 | 5 |
| 15 | EnhBiv | Bivalent Enhancer | 0.63 | 4 |
| 16 | ReprPC | Repressed PolyComb | 1 | 4 |
| 17 | ReprPCWk | Weak Repressed PolyComb | 1 | 4 |
| 18 | Quies | Quiescent/Low | 0.88 | 3 |

*Top value for the Pearson correlation coefficient between each state (18 states) in NIHR ROADMAP et al. (2015) model and our model (12 states). States with correlation coefficients below 0.75 are highlighted in red.

**Table S8. Overview of the genomic and functional features of the 17 regions enriched in macrophage enhancers and cardiovascular disease-associated SNPs.** **The table includes data on region coordinates, enhancer activity profiles, SNPs, GWAS traits, genes in the region, and eGenes identified through macrophage eQTLs.** Abbreviations: Chr. = chromosome, Enh. = enhancer.

|  | **Chr** | **Enh. Start** | **Enh. End** | **Enh. Size (bp)** | **SNPs** | **GWAS trait** | **Enh. Activity** | **Genes in region** | **Gene Type** | **e-Genes** |
| --- | --- | --- | --- | --- | --- | --- | --- | --- | --- | --- |
| 1 | chr1 | 56492700 | 56502700 | 10000 | rs17114036 | Coronary artery disease | M2 | *RP1-158P9.3, PLPP3* | protein_coding | *PLPP3* |
| 2 | chr2 | 64974400 | 64984800 | 10400 | rs10211524 | Metabolite levels (Val) | M1\|M0, M1 | *LINC02245* | lncRNA | *--* |
| 3 | chr2 | 44960600 | 44970600 | 10000 | rs895636 | Blood glucose levels | M1\|M0 | *AC012354.6* | lncRNA | *SRBD1, PPM1B, PREPL, SIX3, LRPPRC, CAMKMT, PRKCE* |
| 4 | chr5 | 173933700 | 173943700 | 10000 | rs6861681 | Waist-hip ratio | M2 | *CPEB4* | protein_coding | *CPEB4* |
| 5 | chr6 | 6731700 | 6743900 | 12200 | rs1294410, rs1294421 | Waist-hip ratio | M2 | *RP1-80N2.2, RP3-470L22.2* | lncRNA | *RP1-80N2.2* |
| 6 | chr8 | 20005100 | 20017000 | 11900 | rs1441756, rs2083637 | Metabolic syndrome (bivariate traits), HDL cholesterol | M1, M1\|M0, M1\|M0 | *--* | -- | *PSD3, LPL* |
| 7 | chr8 | 19952400 | 19978600 | 26200 | rs295, rs301, rs264, rs326, rs331, rs325, rs1059611, rs15285, rs13702, rs2197089, rs10096633, rs10105606, rs17482753 | Metabolic syndrome, Metabolic syndrome (bivariate traits), Coronary artery disease, HDL cholesterol, Triglycerides, Lipoprotein particle size, HDL cholesterol, Triglycerides-Blood Pressure (TG-BP), HDL Cholesterol - Triglycerides (HDLC-TG), Triglycerides | M2, M2\|M1, M2, M2\|M1\|M0, M1, M1\|M0 | *LPL* | protein_coding, -- | *LPL* |
| 8 | chr9 | 104823200 | 104836100 | 12900 | rs4149310, rs2515629 | Large HDL particle size, HDL cholesterol | M1, M1\|M0 | *ABCA1* | protein_coding | *ABCA1* |
| 9 | chr10 | 31699600 | 31709600 | 10000 | rs7081678 | Waist-hip ratio | M1 | *MACORIS* | lncRNA | *MACORIS* |
| 10 | chr11 | 47250200 | 47269900 | 19700 | rs10838681, rs7120118 | Metabolic syndrome, HDL cholesterol | M1\|M0, M2\|M0, M2 | *NR1H3, MADD* | protein_coding | *NR1H3, MADD* |
| 11 | chr12 | 26294000 | 26304000 | 10000 | rs718314 | Waist-hip ratio | M1 | *SSPN, RP11-283G6.4, RP11-283G6.5, RP11-612B6.6* | protein_coding, lncRNA | *SSPN, RP11-283G6.4* |
| 12 | chr15 | 58379600 | 58392000 | 12400 | rs10468017, rs2043085, rs1532085 | Metabolic syndrome (bivariate traits), HDL cholesterol, Triglycerides, Cholesterol Total | M1 | *ALDH1A2* | protein_coding | *ALDH1A2* |
| 13 | chr15 | 59192600 | 59205300 | 12700 | rs2306786 | Triglycerides | M1, M1\|M0 | *MYO1E* | protein_coding | *MYO1E* |
| 14 | chr19 | 44889700 | 44924300 | 34600 | rs157580, rs157582, rs439401, rs445925, rs12721054 | HDL cholesterol, Metabolic syndrome, HDL Cholesterol - Triglycerides (HDLC-TG), Triglycerides, Atherosclerosis, Triglycerides | M1\|M0, M2\|M0, M1, M2\|M1, M1 | *CTB-129P6.4, TOMM40, APOE, CTB-129P6.7, APOC1* | lncRNA, protein_coding, protein_coding, TEC, TEC, protein_coding | *TOMM40, APOE, APOC1* |
| 15 | chr20 | 45908200 | 45919400 | 11200 | rs4810479 | HDL cholesterol, Triglycerides | M1 | *PLTP* | protein_coding | *PLTP* |
| 16 | chr22 | 43923400 | 43933400 | 10000 | rs12483959 | Triglycerides | M2 | *PNPLA3* | protein_coding | *PNPLA3* |
| 17 | chrX | 67719600 | 67729600 | 10000 | rs5031002 | LDL cholesterol | M1 | *AR* | protein_coding | *OPHN1, STARD8, YIPF6* |

**Table S9. Expression patterns of potential enhancer gene targets enriched in CVD-associated regions according to Human Protein Atlas data**

| **Gene** | **Tissue expression cluster** | **RNA tissue cell type enrichment** | **RNA single cell type specific nTPM** |
| --- | --- | --- | --- |
| *PLPP3* | Cluster 89: Fibroblasts - ECM organization | Kidney - Endothelial cells, Prostate - Fibroblasts, Skin - Fibroblast_2, Thyroid - Thyroid glandular cells | Astrocytes: 362.5; Fibroblasts: 403.1; Leydig cells: 881.4; Peritubular cells: 311.7 |
| *LPL* | Cluster 82: **Adipose tissue** - ECM organization | Adipose subcutaneous - Adipocytes (Subcutaneous), Adipose visceral - Adipocytes (Visceral), Breast - Adipocytes (Breast), Skin - Adipocytes (Skin), Testis - Endothelial cells | Adipocytes: 409.8; Cardiomyocytes: 664.2; Granulosa cells: 513.8; Schwann cells: 156.9 |
| *NR1H3* | Cluster 64: **Macrophages** - Immune response | Breast - Adipocytes (Breast), Testis - Early spermatids, Testis - Late spermatids | Hepatocytes: 58.9; **Hofbauer cells**: 47.6; Late spermatids: 48.2; Proximal enterocytes: 68.3 |
| *PLTP* | Cluster 82: **Adipose tissue** - ECM organization | **Adipose subcutaneous - Macrophages, Adipose visceral - Macrophages, Colon - Macrophages**, Lung - Fibroblast_2, Prostate - Fibroblasts, **Skeletal muscle - Macrophages, Thyroid - Macrophages** | **Hofbauer cells**: 1473.5 |
| *APOC1* | Cluster 62: **Liver** - Hemostasis | Liver - Hepatocytes, **Lung - Macrophages**, Skin - Sebaceous gland cells | Hepatocytes: 34731.7 |
| *ALDH1A2* | Cluster 89: Fibroblasts - ECM organization | Adipose visceral - Mesothelial cells, Heart muscle - Fibroblasts, Skeletal muscle - Fibroblasts, Testis - Early spermatids | Early spermatids: 122.1; Endometrial stromal cells: 137.8; Late spermatids: 46.6; **Microglial cells**: 41.0; Spermatocytes: 62.2; Thymic epithelial cells: 82.1 |
| *ABCA1* | Cluster 15: **Liver** - Metabolism | Breast - Adipocytes (Breast) | granulocytes: 120.9; Hepatocytes: 136.6; **Langerhans cells**: 127.5; **Macrophages**: 110.5 |
| *PNPLA3* | Cluster 85: **Liver** - Metabolism | Adipose visceral - Adipocytes (Visceral), Kidney - Proximal tubular cells, Kidney - Proximal tubular cells, Skin - Keratinocyte (granular) | Bipolar cells: 33.8; Hepatocytes: 36.1; Rod photoreceptor cells: 35.9 |
| *APOE* | Cluster 15: **Liver** - Metabolism | Heart muscle - Fibroblasts, Kidney - Proximal tubular cells, Kidney - Proximal tubular cells, Liver - Hepatocytes, **Lung - Macrophages, Pancreas - Macrophages**, Skeletal muscle - Fibroblasts | Hepatocytes: 8379.2; **Hofbauer cells**: 5636.7; Leydig cells: 2655.2; Melanocytes: 3312.6; Muller glia cells: 8525.9; Peritubular cells: 4801.9; Proximal tubular cells: 6176.1; Theca cells: 8535.1 |
| *STARD8* | Cluster 7: **Adipose tissue** - Mixed function | Breast - Endothelial cells, Testis - Endothelial cells | Adipocytes: 23.9; Endothelial cells: 24.8; **Kupffer cells**: 10.2; **Langerhans cells**: 17.0; **Macrophages**: 14.6; monocytes: 29.5; Schwann cells: 17.6 |
| *CPEB4* | Cluster 56: Non-specific - Unknown function | NA | Oligodendrocyte precursor cells: 302.3 |

Table S10. eQTL data from naïve macrophages linking non-coding CVD risk variants with potential enhancer gene targets.

| **rsid** | **Gene name** | **Gene type** | **Data Sources** | **GWAS trait** | **ref** | **alt** | **variant** |
| --- | --- | --- | --- | --- | --- | --- | --- |
| rs2515629 | *ABCA1* | protein | EBI_Alasoo_2018, EBI_Nedelec_2016 | HDL cholesterol | A | G | chr9_104832083_A_G |
| rs4149310 | *ABCA1* | protein | EBI_Alasoo_2018, EBI_Nedelec_2016 | Metabolite levels | A | T | chr9_104826853_A_T |
| rs10468017 | *ALDH1A2* | protein | EBI_Alasoo_2018, EBI_Nedelec_2016 | Metabolic syndrome (bivariate traits) | C | T | chr15_58386313_C_T |
| rs1532085 | *ALDH1A2* | protein | EBI_Alasoo_2018, EBI_Nedelec_2016 | Metabolite levels | A | G | chr15_58391167_A_G |
| rs2043085 | *ALDH1A2* | protein | EBI_Alasoo_2018, EBI_Nedelec_2016 | Metabolic syndrome (bivariate traits) | T | C | chr15_58388755_T_C |
| rs2306786 | *ALDH1A2* | protein | EBI_Alasoo_2018, EBI_Nedelec_2016 | Metabolite levels | C | G | chr15_59195731_C_G |
| rs12721054 | *APOC1* | protein | EBI_Nedelec_2016 | Triglycerides | A | G | chr19_44919330_A_G |
| rs157580 | *APOC1* | protein | EBI_Alasoo_2018, EBI_Nedelec_2016 | HDL cholesterol | G | A | chr19_44892009_G_A |
| rs157582 | *APOC1* | protein | EBI_Alasoo_2018, EBI_Nedelec_2016 | Metabolic syndrome | C | T | chr19_44892962_C_T |
| rs439401 | *APOC1* | protein | EBI_Alasoo_2018, EBI_Nedelec_2016 | HDL Cholesterol - Triglycerides (HDLC-TG), Triglycerides | T | C | chr19_44911194_T_C |
| rs445925 | *APOC1* | protein | EBI_Alasoo_2018, EBI_Nedelec_2016 | Metabolite levels | G | A | chr19_44912383_G_A |
| rs12721054 | *APOE* | protein | EBI_Nedelec_2016 | Triglycerides | A | G | chr19_44919330_A_G |
| rs157580 | *APOE* | protein | EBI_Alasoo_2018, EBI_Nedelec_2016 | HDL cholesterol | G | A | chr19_44892009_G_A |
| rs157582 | *APOE* | protein | EBI_Alasoo_2018, EBI_Nedelec_2016 | Metabolic syndrome | C | T | chr19_44892962_C_T |
| rs439401 | *APOE* | protein | EBI_Alasoo_2018, EBI_Nedelec_2016 | HDL Cholesterol - Triglycerides (HDLC-TG), Triglycerides | T | C | chr19_44911194_T_C |
| rs445925 | *APOE* | protein | EBI_Alasoo_2018, EBI_Nedelec_2016 | Metabolite levels | G | A | chr19_44912383_G_A |
| rs6861681 | *CPEB4* | protein | EBI_Alasoo_2018, EBI_Nedelec_2016 | Waist-hip ratio | G | A | chr5_173935455_G_A |
| rs12721054 | *CTB-129P6.4* | lncRNA | EBI_Nedelec_2016 | Triglycerides | A | G | chr19_44919330_A_G |
| rs157580 | *CTB-129P6.4* | lncRNA | EBI_Alasoo_2018, EBI_Nedelec_2016 | HDL cholesterol | G | A | chr19_44892009_G_A |
| rs157582 | *CTB-129P6.4* | lncRNA | EBI_Alasoo_2018, EBI_Nedelec_2016 | Metabolic syndrome | C | T | chr19_44892962_C_T |
| rs439401 | *CTB-129P6.4* | lncRNA | EBI_Alasoo_2018, EBI_Nedelec_2016 | HDL Cholesterol - Triglycerides (HDLC-TG), Triglycerides | T | C | chr19_44911194_T_C |
| rs445925 | *CTB-129P6.4* | lncRNA | EBI_Alasoo_2018, EBI_Nedelec_2016 | Metabolite levels | G | A | chr19_44912383_G_A |
| rs10096633 | *LPL* | protein | EBI_Alasoo_2018, EBI_Nedelec_2016 | HDL cholesterol, Metabolic traits, Triglycerides | C | T | chr8_19973410_C_T |
| rs10105606 | *LPL* | protein | EBI_Alasoo_2018, EBI_Nedelec_2016 | Triglycerides | C | A | chr8_19970337_C_A |
| rs1059611 | *LPL* | protein | EBI_Alasoo_2018, EBI_Nedelec_2016 | Lipid metabolism phenotypes | T | C | chr8_19967052_T_C |
| rs13702 | *LPL* | protein | EBI_Alasoo_2018, EBI_Nedelec_2016 | HDL Cholesterol - Triglycerides (HDLC-TG) | T | C | chr8_19966981_T_C |
| rs1441756 | *LPL* | protein | EBI_Alasoo_2018, EBI_Nedelec_2016 | Metabolic syndrome (bivariate traits) | A | C | chr8_20010875_A_C |
| rs15285 | *LPL* | protein | EBI_Alasoo_2018, EBI_Nedelec_2016 | Triglycerides-Blood Pressure (TG-BP) | C | T | chr8_19967156_C_T |
| rs17482753 | *LPL* | protein | EBI_Alasoo_2018, EBI_Nedelec_2016 | HDL cholesterol | G | T | chr8_19975135_G_T |
| rs2083637 | *LPL* | protein | EBI_Alasoo_2018, EBI_Nedelec_2016 | HDL cholesterol | A | G | chr8_20007664_A_G |
| rs2197089 | *LPL* | protein | EBI_Alasoo_2018, EBI_Nedelec_2016 | Metabolic syndrome (bivariate traits) | G | A | chr8_19968862_G_A |
| rs264 | *LPL* | protein | EBI_Alasoo_2018, EBI_Nedelec_2016 | Coronary artery disease | G | A | chr8_19955669_G_A |
| rs295 | *LPL* | protein | EBI_Alasoo_2018, EBI_Nedelec_2016 | Metabolic syndrome | A | C | chr8_19958727_A_C |
| rs301 | *LPL* | protein | EBI_Alasoo_2018, EBI_Nedelec_2016 | Metabolic syndrome (bivariate traits) | T | C | chr8_19959423_T_C |
| rs325 | *LPL* | protein | EBI_Alasoo_2018, EBI_Nedelec_2016 | HDL cholesterol | T | C | chr8_19961817_T_C |
| rs326 | *LPL* | protein | EBI_Alasoo_2018, EBI_Nedelec_2016 | HDL cholesterol, Triglycerides | A | G | chr8_19961928_A_G |
| rs331 | *LPL* | protein | EBI_Alasoo_2018, EBI_Nedelec_2016 | Lipid metabolism phenotypes | G | A | chr8_19962894_G_A |
| rs7081678 | *MACORIS* | lncRNA | EBI_Alasoo_2018, EBI_Nedelec_2016 | Waist-hip ratio | G | A | chr10_31701695_G_A |
| rs10838681 | *MADD* | protein | EBI_Alasoo_2018, EBI_Nedelec_2016 | Metabolic syndrome | G | A | chr11_47253513_G_A |
| rs7120118 | *MADD* | protein | EBI_Alasoo_2018, EBI_Nedelec_2016 | HDL cholesterol | T | C | chr11_47264739_T_C |
| rs10468017 | *MYO1E* | protein | EBI_Alasoo_2018, EBI_Nedelec_2016 | Metabolic syndrome (bivariate traits) | C | T | chr15_58386313_C_T |
| rs1532085 | *MYO1E* | protein | EBI_Alasoo_2018, EBI_Nedelec_2016 | Metabolite levels | A | G | chr15_58391167_A_G |
| rs2043085 | *MYO1E* | protein | EBI_Alasoo_2018, EBI_Nedelec_2016 | Metabolic syndrome (bivariate traits) | T | C | chr15_58388755_T_C |
| rs2306786 | *MYO1E* | protein | EBI_Alasoo_2018, EBI_Nedelec_2016 | Metabolite levels | C | G | chr15_59195731_C_G |
| rs10838681 | *NR1H3* | protein | EBI_Alasoo_2018, EBI_Nedelec_2016 | Metabolic syndrome | G | A | chr11_47253513_G_A |
| rs7120118 | *NR1H3* | protein | EBI_Alasoo_2018, EBI_Nedelec_2016 | HDL cholesterol | T | C | chr11_47264739_T_C |
| rs17114036 | *PLPP3* | protein | EBI_Alasoo_2018, EBI_Nedelec_2016 | Coronary artery disease | A | G | chr1_56497149_A_G |
| rs4810479 | *PLTP* | protein | EBI_Alasoo_2018, EBI_Nedelec_2016 | Metabolite levels | C | T | chr20_45916409_C_T |
| rs12483959 | *PNPLA3* | protein | EBI_Alasoo_2018 | Metabolite levels | G | A | chr22_43930116_G_A |
| rs718314 | *RP11-283G6.4* | lncRNA | EBI_Alasoo_2018 | Waist-hip ratio | A | G | chr12_26300350_A_G |
| rs1294410 | *RP1-80N2.2* | lncRNA | EBI_Alasoo_2018, EBI_Nedelec_2016 | Waist-hip ratio | T | C | chr6_6738519_T_C |
| rs1294421 | *RP1-80N2.2* | lncRNA | EBI_Alasoo_2018, EBI_Nedelec_2016 | Waist-hip ratio | T | G | chr6_6742916_T_G |
| rs718314 | *SSPN* | protein | EBI_Alasoo_2018 | Waist-hip ratio | A | G | chr12_26300350_A_G |
| rs12721054 | *TOMM40* | protein | EBI_Nedelec_2016 | Triglycerides | A | G | chr19_44919330_A_G |
| rs157580 | *TOMM40* | protein | EBI_Alasoo_2018, EBI_Nedelec_2016 | HDL cholesterol | G | A | chr19_44892009_G_A |
| rs157582 | *TOMM40* | protein | EBI_Alasoo_2018, EBI_Nedelec_2016 | Metabolic syndrome | C | T | chr19_44892962_C_T |
| rs439401 | *TOMM40* | protein | EBI_Alasoo_2018, EBI_Nedelec_2016 | HDL Cholesterol - Triglycerides (HDLC-TG), Triglycerides | T | C | chr19_44911194_T_C |
| rs445925 | *TOMM40* | protein | EBI_Alasoo_2018, EBI_Nedelec_2016 | Metabolite levels | G | A | chr19_44912383_G_A |
